# Breaking free from the clock’s tyranny restores memory to brain damaged flies

**DOI:** 10.1101/2024.01.25.577231

**Authors:** Stephane Dissel, Ellen Morgan, Lijuan Cao, Zachary Peters Wakefield, Shohan Shetty, Dorothy Chan, Vincent Duong, Jeff Donlea, Hamza Farah, Vasilios Loutrianakis, Melanie Ford, Lillith Streett, Erica Periandri, Zhaoyi Li, Irene Huang, Dina Abdala, Arjan Kalra, Lea Sousani, Brandon Holder, Chloe McAdams, Bruno van Swinderen, Paul J. Shaw

## Abstract

The relationship between sleep and memory is an active topic of investigation. In this context, we demonstrate that enhancing sleep restores memory to flies with ablated Mushroom Bodies (MB), a key memory center; this is consistent across several memory assays. Mapping the underlying circuitry reveals circadian modulation of a subset of Dopaminergic neurons (DANs) that modulate aversive learning. Using imaging, we show that MB-ablation disrupts, and sleep restores the time of day these neurons are most responsive. Knocking down the receptor for the clock output signal, *Pigment-dispersing factor* (Pdfr), in this subset of DANs restores memory to MB-ablated flies. Crucially, MB-ablation does not result in memory impairments in the absence of a functioning clock. Our results reveal neuromodulation’s key role in cognitive restoration, where sleep aids memory in damaged brains, but a functioning clock unexpectedly hinders this process.

## Introduction

Recent studies have demonstrated that enhancing sleep can restore memory to flies with mutations in classic memory genes and to flies expressing human Alzheimer’s genes (*1–7*). Specifically, enhancing sleep for 2 days in mutants for the *Drosophila adenyl cyclase*, *rutabaga* (*rut^2080^*), restores memory when assessed by Aversive Phototaxic Suppression (APS), Courtship Conditioning, Place Learning and Space Learning (*1, 2, 4*). Each of these memory assays utilizes different stimuli and relies upon different neuronal circuits, transmitters, and peptides (*8–10*). An important, and often overlooked feature of the sleep induction protocol, is that sleep restores brain functionality to animals before they are exposed to the training/testing conditions (*4*). In each case, the fly has no way of knowing or anticipating that it will eventually be exposed to a particular learning environment or how. Interestingly, when sleep induction protocols are terminated, flies slowly revert back to being memory impaired over the course of 72 h (*4*). These data indicate that experimentally increasing sleep actively alters and maintains the integrity of neural circuits supporting adaptive behavior.

The molecular mechanisms recruited by sleep to restore memory to impaired flies remains elusive. However, the literature suggests a potential role for sleep in recruiting broader neuromodulatory pathways. These pathways integrate environmental context and internal state, thereby altering the configuration of neural circuits to enhance behavioral responses and adaptability in specific contexts (*11–15*). Dopaminergic neurons play an important and evolutionarily conserved role for providing information about rewards and punishments and for reconfiguring neuronal circuits according to internal state (*16–23*). Interestingly, while the activity of dopaminergic neurons has a defined impact on downstream memory circuits in response to the unconditioned stimulus, dopamine (DA) also interacts with other circuits, including the circadian clock, to induce long-lasting modulation of neural networks regulating motivated behavior (*23–31*). Given that the time-course for sleep to restore memory to cognitively impaired flies is slow (∼12-24 h), we hypothesize that sleep improves

performance by recruiting neuromodulatory pathways to optimize information processing in the brain. In flies, clock circuits play an active role in memory formation, regulate sleep and waking, and impose daily activity rhythms onto dopaminergic neurons (*32–37*). Thus, here we examine the intricate interplay between sleep, clock circuits, and dopaminergic modulatory pathways in shaping memory formation in flies suffering catastrophic brain damage. We hypothesize that elucidating the mechanisms by which sleep reinstates adaptive behavior in impaired flies will not only shed light on specific aspects of sleep regulation and function but may also uncover novel principles of neuromodulation in general.

## Results

### Enhanced sleep restores short-term memory to Mushroom Body ablated flies

The Mushroom Body (MB) serves as a key center for associative memory, sensory processing and decision making (*38*). Kenyon cells (KCs), the intrinsic neurons of the MB, form three distinct lobes (α/β, α’/ β’ and γ lobes) that synapse onto 21 different types of MB output neurons (MBONs) whose dendrites project to defined compartments within the MB neuropils (*38*). The interactions between KCs and MBONs are modulated by 20 distinct populations of dopaminergic neurons (DANs) which convey reward or punishment.

The importance of the MB in associative olfactory memory was first confirmed using a chemical ablation method that kills KCs (*39*). MB ablation was later used to demonstrate the importance of the MB for courtship conditioning and Aversive Phototaxic Suppression (APS) (*40, 41*). Thus, as a stringent test of sleep’s capacity to restore function in significantly compromised brains, we intensified sleep in flies with MB ablations and analyzed Short-Term Memory (STM) using the APS, and Long-Term Memory (LTM) using courtship conditioning. To enhance sleep, we employed two independent and widely utilized approaches: Gaboxadol administration (*4, 42–55*), and genetically activating sleep promoting neurons (*56–66*). Newly hatched *Canton-S* (*Cs*) larvae were fed Hydroxyurea (HU) to eliminate the Kenyon cells forming the α/β, α’/β’ and most of the γ lobes (*39*). To confirm the effectiveness of HU treatment, we evaluated sleep and used histology to visualize KCs. Consistent with previous reports (*41, 67, 68*), MB ablation significantly disrupts sleep and effectively eliminates most Kenyon cells (Figure 1A, A’). Importantly, Gaboxadol substantially increased sleep in MB-ablated flies (Figure 1A). MB-ablated flies did not exhibit STM compared to sham-treated controls. However, after two days of enhanced sleep MB-ablated flies displayed an STM when subsequently tested in the APS (Figure 1B) (*41*). Note that individual scores will not be plotted with group means due to the structure of data obtained using the APS (*4, 69, 70*). However, to ensure transparency, individual scores are reported in Supplemental File 1. Evaluation of photosensitivity (PSI) and quinine sensitivity (QSI) (Figure 1B’) confirmed that sensory processes were unaffected by MB ablation or Gaboxadol, consistent with previous findings (*4, 41*). Therefore, Gaboxadol-induced sleep can restore STM in MB-ablated flies.

**Figure 1.**
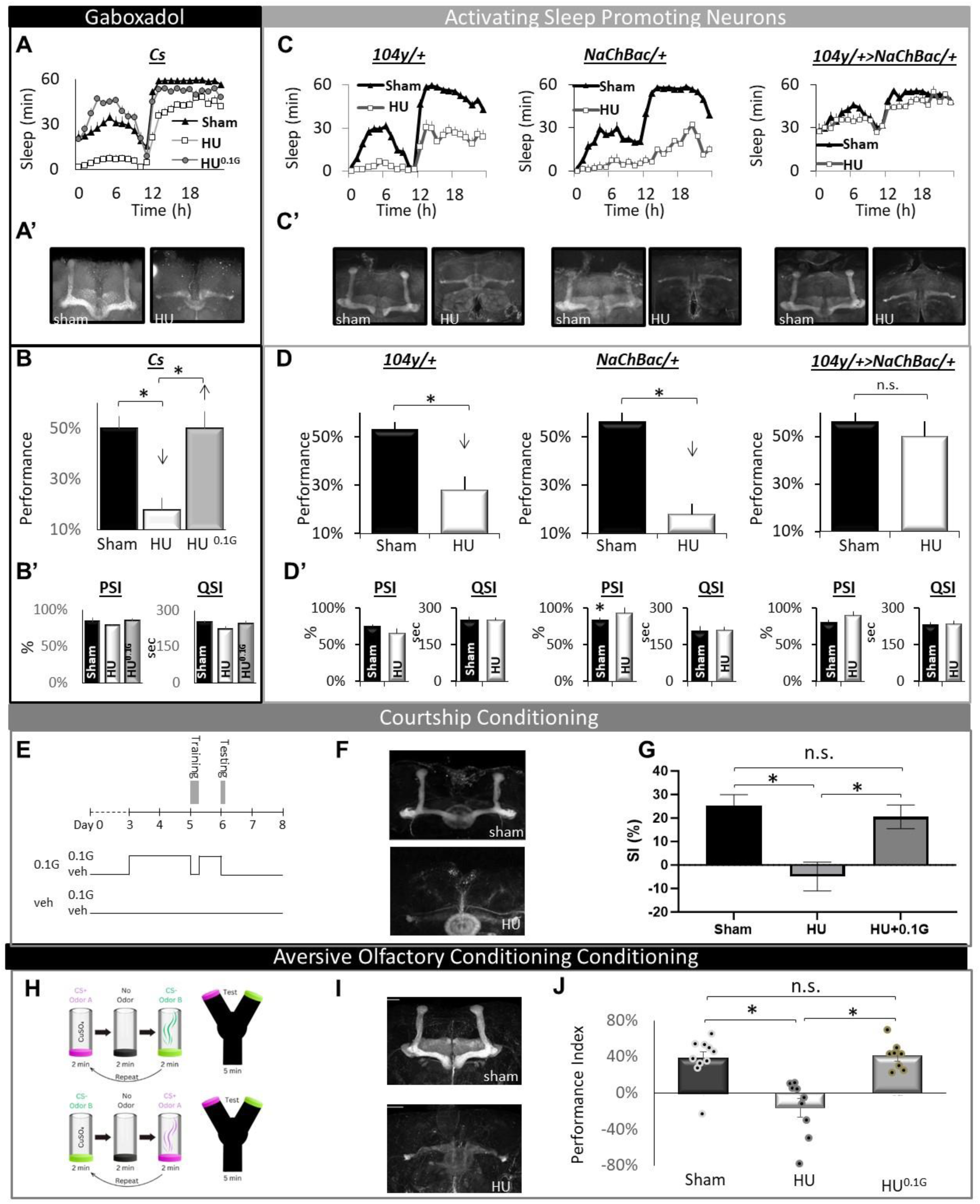
Sleep restores memory to Mushroom Body ablated flies. **(A)** Sleep is reduced in Hydroxy Urea (HU)-fed, MB-ablated flies (HU, white, n=26) flies compared to sham treated (black, n=29) , and age-matched HU ablated siblings fed 01.mg/mL (HU^0.1G^, grey, n=18) (See Table 1 for statistics)**. (A’)** Whole-mount anti-Fasciclin2 (Fas2) immunohistochemistry verifying MB-ablation. B) MB-ablated *Cs* flies exhibit impaired STM (black) compared to either sham treated siblings or Gaboxadol-fed, HU-ablated flies display wild-type STM, *p<0.05 modified Bonferroni test. **(B’)** Neither HU nor Gaboxadol altered PSI or QSI, *p<0.05 modified Bonferroni test. **C)** Sleep in minutes/hour of sham-treated (black) and HU-treated (white) flies. Parental controls (*UAS*-*NaChBac*/*+*, and *104y*/*+*) exhibit a significant decrease in sleep after HU treatment compared to sham-treated controls. HU-treatment does not alter sleep in *104y*/*+>UAS*-*NaChBac*/*+* flies (n=12-16 each group). **(C’)** αFas2 immunohistochemistry verifying MB-ablation. **(D)** HU-fed *104y-GAL4/+* and *UAS-NaChBac/+* controls exhibit substantial impairments in STM (white) compared to sham-ablated controls (black). HU-treated *104y-GAL4/+>UAS-NaChBac/+* flies maintain wild-type STM,*p<0.05 modified Bonferroni test n=>8/group). **(D’)** Neither genotype nor HU treatment altered PSI or QSI, *p<0.05 modified Bonferroni test. **(E)** Schematic of the spaced training protocol used for Gaboxadol-induced sleep. Naïve male *Cs* flies were fed 0.1 mg/ml Gaboxadol (0.1G) or vehicle (veh) from age day 3 to day 5. Gaboxadol fed flies were removed from Gaboxadol 1 h prior to and during spaced training and then returned to 0.1G for 24 h post-training. Vehicle-fed flies were maintained on vehicle throughout the protocol. Courtship was tested in all groups 48 h after training on day 7. **(F)** Fas2 staining. **(G)** SI (Suppression Index= 100*[1-CI^train+^/CI^train-^]) for vehicle-fed Cs (Sham), HU-ablated Cs (Cs HU) and Gaboxadol-fed HU-ablated Cs (Cs HU+0.1G) flies. LTM is disrupted in HU-ablated Cs flies but it is similar to Sham treated flies in Gaboxadol-fed HU-ablated Cs flies; ,*p<0.05 modified Bonferoni test, n=28-29 flies per condition). **(H)** Schematic of Aversive Olfactory learning. Flies were trained with either Odor A (MCH) or Odor B (OCT) as CS+. During training, flies experienced 80 mM CuSO4 (US) with the CS+. Immediate memory was evaluated by measuring the preference for CS+ and the alternative CS-. The Performance Index (PI) = (CS+ choices - CS- choices)/total flies). Scores from the fly groups trained with OCT or MCH as CS+ and were averaged to derive a data point. **(I)** Fas2 staining. **(J)** Sham-treated Cs flies (n=10) had higher PI than HU-treated siblings (n=9). PI restored in HU-treated flies on Gaboxadol for 2 days (n=8); *p<0.05 modified Bonferroni test. Statistics are reported in Supplemental Table 2.

To confirm that the memory enhancement associated with Gaboxadol is due to sleep rather than nonspecific effects of the drug, we used genetic tools to express the bacterial sodium channel *NaChBac* in the *104y-GAL4*, sleep-promoting neurons (*43, 57, 58, 71, 72*). As seen in Figure 1C, HU treatment substantially decreased sleep in *104y-GAL4/+* and *UAS-NaChBac/+* parental controls compared to sham-treated siblings. Despite eliminating most Kenyon cells (Figure 1C’), sleep was maintained in HU-fed *104y-GAL4/+>UAS-NaChBac/+* experimental flies (Figure 1C). Moreover, STM was impaired in HU-fed *104y-GAL4/+* and *UAS-NaChBac/+* parental controls compared to sham-treated siblings (Figure 1D). In contrast, STM was preserved in HU-fed *104y-GAL4/+>UAS-NaChBac* flies. As above, no changes in PSI and QSI were observed, indicating that none of the changes in STM are due to changes in sensory thresholds (Figure 1D’). Thus, increasing sleep using two independent methods, feeding flies Gaboxadol or genetically activating sleep-promoting neurons, can restore STM to MB-ablated flies.

### Enhanced sleep restores long-term memory to Mushroom Body ablated flies

We have previously used courtship conditioning to show that enhanced sleep can restore LTM to the canonical memory mutant *rutabaga* (*rut^2080^*) (*4*). In courtship conditioning, males whose courtship attempts are rejected by recently mated females form long-term associative memories as evidenced by subsequently reduced courtship of a receptive virgin female (*58, 73, 74*). A previously validated protocol for increasing sleep during courtship training is shown in Figure 1E (*4*). Courtship conditioning was performed on 1) untreated *Cs* controls, 2) HU-treated siblings, and 3) HU-treated Gaboxadol-fed siblings. HU effectively eliminates most Kenyon cells (Figure 1F). As seen in Figure 1G, space training in control *Cs* flies exhibits LTM as defined by courtship suppression. Not surprisingly, spaced training did not induce LTM in MB-ablated flies (Figure 1G) (*40*). However, MB-ablated flies with enhanced sleep formed a LTM (Figure 1G). These data indicate that the ability of sleep to restore function in severely impaired brains is not confined to a single memory assay or memory type.

### Enhanced sleep restores aversive olfactory memory to Mushroom Body ablated flies

To further examine the role of sleep in restoring STM to brain damaged flies, we fed larvae HU and examined aversive olfactory memory using the recently described Y-maze (*75*) (Figure 1H). As above, HU was effective in eliminating most KCs (Figure 1I). Importantly, Vehicle-fed controls display an intact immediate memory while, MB-ablated flies are significantly impaired (Figure 1J), as expected (*39*). In contrast, Gaboxadol-fed, MB-ablated siblings display intact STM (Figure 1J). Neither HU nor Gaboxadol altered odor acuity (Supplemental Table 1). Thus, Gaboxadol restores STM to MB-ablated flies when using three independent memory assays.

### Enhanced Sleep can restore STM to canonical memory mutants with disordered KCs

The APS has demonstrated remarkable effectiveness in reproducing results obtained using aversive olfactory conditioning, including for mutants which disrupt MB morphology (*41, 70, 76–79*). Given that the APS is a highly sensitive assay for investigating the relationship between sleep and plasticity, we will focus on the APS for the remaining studies (*3, 80–82*). To extend our results beyond chemically ablating the MBs, we examined the effect of Gaboxadol-induced sleep on STM in fly’s mutant for *linotte/derailed* (*lio^1^* or *lio^2^*), *Origin recognition complex subunit 3* (*lat^6^*) and *mushroom bodies undersized* (*w; CK2β^mbu^/+*). As seen in Supplemental Figure S1 A-L, STM is impaired in *lio^1^*, *lio^2^*, *lat^6^* and *CK2β^mbu^*mutants as expected. STM impairments are reversed following 48 h of Gaboxadol induced sleep (Supplemental Figure S1A-L). Notably, *lio^1^*, *lio^2^* and *lat^6^*mutants sleep deprived while on Gaboxadol did not exhibit STM, indicating that the improved STM was not due to off-target effects of Gaboxadol, consistent with previous reports (Supplemental Figure S1A, B) (*4, 83*). We also used a within-subject design to verify that Gaboxadol-induced sleep could restore STM to individual *lio^2^* mutant flies and verified STM impairments in *lat^6^* mutants using RNAi (Supplemental Figure S1D-L). Thus, sleep can restore STM to genetic mutants with disruptions in MB morphology.

To test whether sleep could restore STM by modulating MB neurons that might have survived HU ablation, we expressed a cold-sensitive mutation of the *ricin-toxin A chain* (*UAS-Ricin^CS^*), which blocks protein synthesis, using a pan-MB driver (*247-GAL4/+*> *UAS-Ricin^CS^*/+) or a driver that is strongly expressed in the γ lobes (*201y- GAL4/+> UAS-Ricin^CS^*/+) (*84*). Flies were reared at 19°C and then kept at 30°C for 4 days to block protein synthesis (*84*). Blocking protein synthesis in the MBs for 2 days resulted in STM impairments (data not shown). Thus, to evaluate the effectiveness of Gaboxadol-induced sleep, *247/+>UAS-Ricin^CS^/+* and *201y/+>UAS-Ricin^CS^/+* flies were kept at 30°C on normal food for 2 days to ensure that the MBs would be disrupted (Supplemental Figure S1G). On the morning of day 6, flies were placed onto 0.1mg/mL of Gaboxadol and kept at 30°C for two additional days. Consequently, Gaboxadol-induced sleep was only increased after flies exhibited deficits in STM. Consistent with the data presented above, STM was restored following 2 days of Gaboxadol-induced sleep (Supplemental Figure S1H). Collectively, these findings demonstrate that sleep has the capacity to restore STM in flies with catastrophic disruption of Kenyon cells, even when the disruption arises from a variety of different causes.

### Time of day regulation of STM

How might sleep restore memory to flies with such catastrophic lesions? We hypothesized that without proper instructive signals from the Kenyon cells, the activity of downstream circuits might become increasingly uncoordinated. We further hypothesized that sleep restores balance to neurons that can no longer properly connect to KCs. To test this hypothesis, we evaluated γ-lobe projecting MBONs and DANs. MBON/DAN interactions are critical for memory formation and have been discussed extensively elsewhere (*16, 85–87*). Briefly, activating MBONs innervating the γ1, γ2 and γ3 compartments promotes attraction, while MBONs innervating γ4, and γ5 promote avoidance (*16*). During aversive learning, DA input from PPL1 neurons depress KC→γ1, γ2 synapses thereby reducing their activity and shifting the balance of γ lobe activity in favor of avoidance. During appetitive learning, DA input from PAM neurons depress KC→γ4, γ5 synapses to favor attraction (Supplemental Figure S2A).

To determine whether γ-lobe projecting MBONs participate in the APS, we expressed the *temperature-gated nonspecific cation channel Transient receptor potential cation channel* (*UAS-TrpA1*) in MBONs. The temperature was increased to 30°C overnight to replicate protocols previously used to identify circuits that influence sleep and waking (*16, 88*)(Supplemental Figure S2 A,B). As seen in Supplemental Figure S2C, chronically activating γ-lobe projecting MBONs disrupted STM thereby establishing their role in the APS and emphasizing the harmful consequences of prolonged neural activity in memory circuits.

To further characterize γ lobe-projecting neurons, we used two complementary strategies: a rescue approach to restore functioning to independent γ lobe-projecting MBONs and DANs, and a loss-of-function analysis to disrupt STM. For the rescue experiments, we expressed wild-type *rutabaga* in γ lobe-projecting MBONs and DANs to flies otherwise mutant for *rutabaga* (e.g. *rut^2080^; split-GAL4>UAS-rut^wt^*). This approach has been used to identify KCs that regulate STM (*89, 90*). For the loss-of-function strategy, we knocked down the *Dopamine 1-like receptor 1* (*Dop1R1*) (e.g. *split-GAL4>Dop1R1^RNAi^*), which plays an important role in the APS (*41, 91*).

During our preliminary experiments, we noticed that specific genotypes demonstrated the ability to form STM in the morning whereas their siblings unexpectedly failed to form an STM in the afternoon, and vice versa. Since STM regulation based on the time of day is widespread in the animal kingdom, including *Drosophila* (*31, 92, 93*), we continued by methodically examining STM in the morning (AM) and in the afternoon (PM) for γ lobe-projecting MBONs and DANs and respective controls (Figure 2A-F’ and Supplemental Figure S2, D-K). An example of a compartment that supports the formation of STM in the AM is shown in Figure 2A. Restoring wild-type *rut* in just four γ2α’1-MBONs rescued STM in the AM even though the fly was mutant for *rut^2080^* in all other neurons (*rut^2080^;051B/+* vs. *rut^2080^;051B/+>UAS-rut^WT^*/+; comparisons are between Black bars denoting STM in the AM). Although STM was intact in *rut^2080^; 051B/+>UAS-rut^WT^*/+ in the AM, no STM was observed in siblings tested in the PM (Figure 2A, *rut^2080^;051B/+* vs. *rut^2080^;051B/+>UAS-rut^WT^*/+; comparing between White bars depicting STM in the PM). Similar results were found for two independent γ2-projecting DANs, *060B* (PPL1-γ2,α1, PPL1-α’2α2, PPL1-α’3 and PPL1-α3-projecting DANs) and *296B* (PPL1-γ2,α1 , PPL1-α’2α2) (Figure 2B, C). No STM was observed in *rut^2080^*; *UAS-rut^WT^/+* controls in either the AM or PM (Supplemental Figure S2,D). Interestingly, we observed the opposite result with loss-of-function experiments (Figure 2A’). Specifically, *051B/+>UAS-Dop1R1^RNAi^/+* flies failed to form a STM in the morning (Figure 2A’, *051B/+* vs. *051B/+>UAS-Dop1R1^RNAi^/+*; Black bars), yet their siblings were able to form a STM in the afternoon (Figure 2A’, *051B/+* vs. *051B/+>UAS-Dop1R1^RNAi^/+;* White bars). Crucially, *UAS-Dop1R1^RNAi^/+* parental controls displayed normal STM in both the AM and PM (Supplemental Figure S2H).

**Figure 2.**
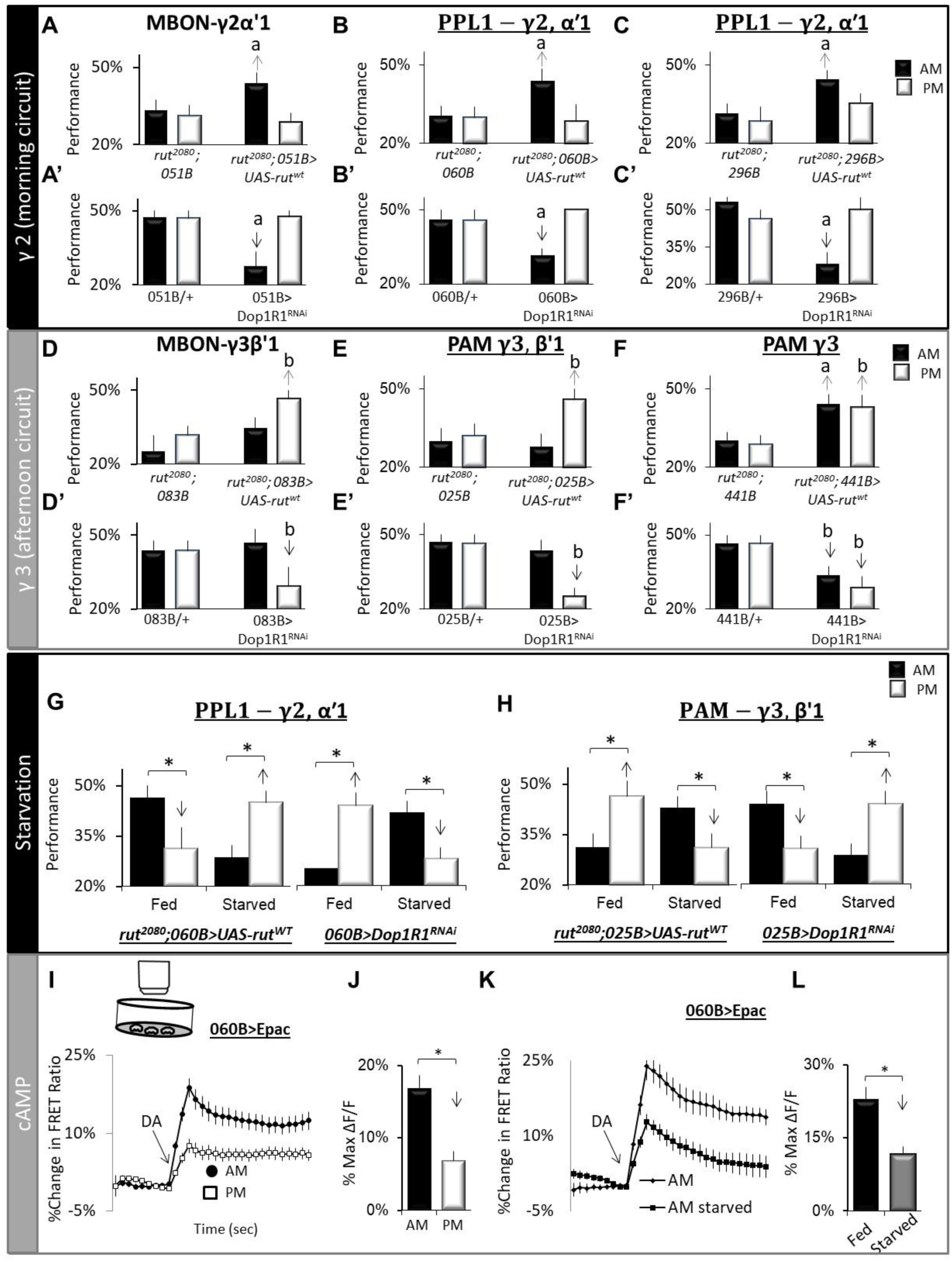
Flies employ distinct strategies for STM at different times of the day. **(A-F)** <u>Rescue:</u> *UAS-rut^WT^* was expressed in a *rut^2080^;Split-GAL4* background and STM was evaluated separately in the morning (AM; 8am to 12pm, black bars) and in the afternoon (PM ; 1pm to 5pm, white bars). Statistical comparison are made between experimental (*rut^2080^; split-GAL4>UAS-rut^WT^* ) and parental controls (*rut^2080^* /+; Split-GAL4/+) for a given time of day (black bars vs black bars or white bars vs white bars; n>8 flies/group; ^a, b^ p<0.05, modified Bonferroni test) (see Table 2 for statistics). **(A’-F’)** <u>Loss-of Function:</u> *UAS-Dop1R1^RNAi^* was expressed using Split-GAL4 drivers and STM was evaluated in the AM and PM. Statistical comparison are made between experimental (*split-GAL4>*D*op1R1^RNAi^*) and parental controls (Split-GAL4/+) for a given time of day (n>8 flies/group; ^a, b^ p<0.05, modified Bonferroni test). **(G)** Rescue and loss of function in paired posterior lateral 1 (PPL1) 060B neurons. In contrast to fed siblings, starved *rut^2080^;060B>UAS-rut^WT^*flies learn in the afternoon but not the morning ; starved loss-of-function *060B>Dop1R1^RNAi^*learn in the morning but not the afternoon, respectively; n>8 flies/group, * p<0.05, modified Bonferroni test) **(H)** Rescue and loss of function in protocerebral anterior medial (PAM), *025B* neurons. Starved *rut^2080^;025B>UAS-rut^WT^* flies learn in the morning but not in the afternoon, while starved *025>Dop1R1^RNAi^* learn in the afternoon but not the morning; n>8 flies/group, * p<0.05, modified Bonferroni test. **(I)** Normalized FRET ratio in *060B-GAL4>UAS-Epac1-camps* in response to DA (3^×^ ^10−5^ ^M^) in the AM (n=22 cells) and PM (n=9). **(J)** Quantification of **I** (ttest p=.0013) **(K)** Normalized FRET ratio in Fed (n=9) and starved (n=18 ) 060B-GAL4>UAS-Epac1-camps in response to DA (3^×^ ^10−5^ ^M^). **(L)** Quantification of **J** (ttest p=.000213). Statistics are reported in Supplemental Table 3.

In contrast to the γ2 compartment, the rescue and loss-of-function studies indicate that γ3-projecting γ3β’1- MBON (083C) and one of its associated DANs (025B, β’1) support STM in the PM but not the AM (Figure 2D,D’ and E,E’). The γ4- and γ5-projecting DANs also support STM in the PM but less so in the AM (Supplemental Figure S2E-J). The ability to form STM in the AM or PM does not appear to be represented at the levels of the KCs (607B, 131B) or the neurons conveying light (R334C-split-GAL4) or bitter information (GR66a-GAL4) as STM in the AM or PM are similarly impacted (Supplemental Figure S2G,K). Interestingly, we identified a GAL4 line (R75H07) that includes neurons downstream of γ2α’1-MBONs. These neurons support STM in the AM but not in the PM, suggesting that temporal information can be transmitted to downstream circuits (Supplemental Figure S2, L-O). Sensory thresholds (PSI and QSI) do not vary over the course of the biological day (*41*) and were not altered in any genotype. Thus, the APS, which requires flies to inhibit their prepotent attraction to light, utilizes both appetitive and aversive γ-lobe projecting MBONs and DANs to regulate STM. Furthermore, these findings suggest that circadian circuits reconfigure the MBON network, enabling flies to employ distinct strategies for STM at different times of the day.

### Starvation alters time of day regulation of STM

Internal states like hunger, thirst, and arousal reconfigure neural circuits, to enable individuals to seamlessly adapt their behavior in response to the fluctuating demands of their environment. One compelling example of this dynamic adaptation is starvation. Starvation fine-tunes the activity of several classes of MBONs and DANs, including those targeting the γ2α’1-compartment and thereby shifts the balance between approach and avoidance pathways (*94, 95*). To determine how time-of-day regulation and motivational state interact to modulate STM, we replicated the rescue and loss-of-function experiments for PPL1-γ2,α1 and PAM-γ3,β’1 DANs in flies maintained on their normal diet (fed) and compared the results to their starved siblings (Figure 1B,B’ E,E’). As seen in Figure 1G, we replicated the morning-STM phenotypes from Figure 1B,B’. However, starved, rescue-siblings no longer displayed a STM in the AM but in the afternoon instead (Figure 2G); the opposite was true for loss-of-function siblings. We also replicated the PM phenotype for fed PAM- γ3,β’1 DANs (Figure 1H). As with the PPL1-γ2,α1 DANs, starvation shifted the time of day the circuit was required for STM (Figure 2H) . These data indicate that while motivational state can alter MB circuits, the segregation of circuits based upon time of day persists.

### Starvation alters the time of day when γ2-projecting DANs are responsive

Given that γ2,α1-projecting MBONs regulate both sleep and STM, we focused on the γ2 compartment for the remaining studies. As noted, the clock imposes daily activity rhythms onto DANs (*36*). Thus, we hypothesized 060B, PPL1-γ2,α1, PPL1-α’2α2, PPL1-α’3 and PPL1-α3-projecting DANs may display changes in activity in the AM vs. PM. To test this hypothesis, we expressed the *cyclic adenosine monophosphate* (cAMP) sensor, *UAS- Epac1-camps*, in *060B* neurons and used high throughput live brain imaging to monitor neuronal responses to bath-applied DA in the AM or PM from cell bodies (Figure 2I) (*14, 96, 97*). As seen in Figure 2I-J, *060B* cell bodies showed robust increases in FRET signaling in the AM compared to the PM. To determine how starvation might influence this relationship, flies were starved and changes in FRET were examined in the AM. As seen in Figure 1K-L, starvation reduced FRET responses in the AM compared to fed siblings, consistent with the behavioral results. To better understand which dopaminergic receptors are mediating time of day effects, we evaluated STM in the AM and PM after knocking down Dop*1R2*, *DopEcR*, or *D2R* receptor in sleep-promoting Kenyon cells projecting to the γ lobe (607B), and in 060B and 051B neurons (Supplemental Figure S2, O,P). Importantly, these data highlight that the γ lobe-projecting MBONs and DANs are under significant modulatory control independent from their role in relaying information about rewards and punishments. Therefore, the presence or absence of STM in a fly is not solely determined by the manipulation of a specific gene but is also influenced by the time of day.

### MB ablation alters the response properties of large ventrolateral neurons LNvs (lLNvs)

Despite memory impairments being the most discussed aspect of MB ablation, flies lacking MBs exhibit prolonged and dramatic increases in waking (*67, 68*). Moreover, sustained MB activity or inactivity impacts circadian locomotor rhythms (*98, 99*). A previous report suggests only modest effects of MB ablation on the clock circuitry (*100*). Nonetheless, we reexamined locomotor activity of MB-ablated flies. We confirmed that HU successfully eliminated KCs (Figure 3A). Moreover, MB ablation reduced the percentage of flies that are rhythmic but did not alter period length (Figure 3B-D). Both morning and evening anticipation were disrupted by MB ablation on day two in both Dark:Dark (DD) and in a 12:12 Light:Dark (LD) schedule (Figure 3E and data not shown) (*101, 102*). Importantly, MB ablation disrupts structural plasticity in the projections of small ventrolateral clock neurons (sLNvs) (Supplemental Figure S3A), and degrades social enrichment, a form of plasticity mediated by the ventrolateral clock neurons (LNvs) (Supplemental Figure S3B) (*103*). Consistent with previous results, live brain imaging of *Pdf-GAL4>UAS-Epac-camps1* flies confirmed that large ventrolateral (lLNvs) respond to DA similarly in the AM and PM (Supplemental Figure S3C) (*96*). However, after MB ablation, *Pdf-GAL4>UAS-Epac-camps1* flies exhibited increased cAMP in response to DA, specifically in the PM (Supplemental Figure S3C). These data suggest that MB ablation may alter aspects of neuromodulation that impact more distal circuits, including the clock circuitry.

**Figure 3:**
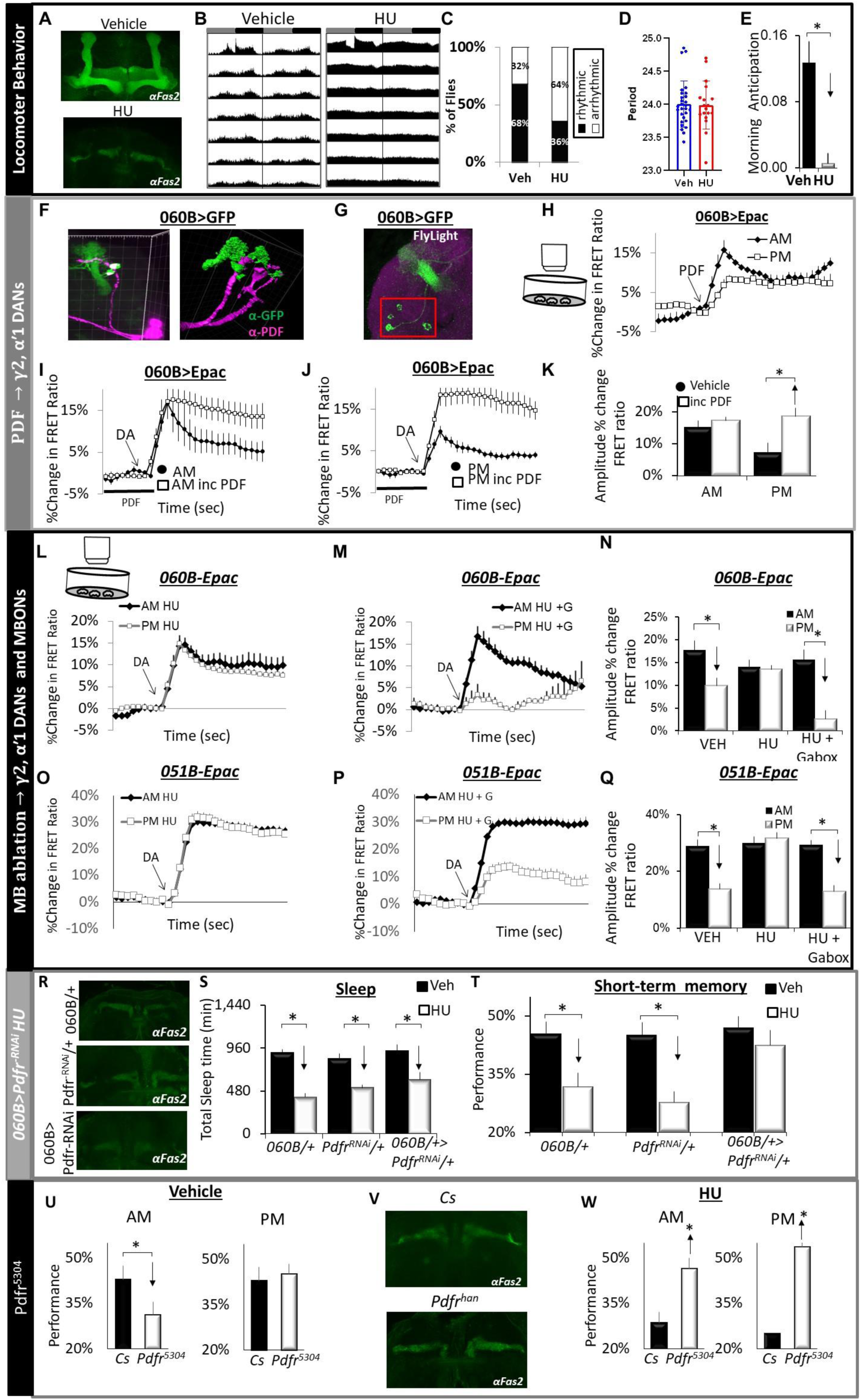
Knocking down Pdfr in ∼8 γ2α’1 projecting DANs restores STM to MB ablated flies. **(A)** Fas2 staining in vehicle and HU fed flies. **(B)** Actograms showing average activity of Cs flies in constant darkness; data is double-plotted. Dark gray rectangles = subjective day, black rectangles (n=40-50 flies/condition) **C)** HU reduces the % of rhythmic flies; chi-square =9.39; p =0.002. **(D)** Scatter plot showing the period of rhythmic flies. **(E)** Morning anticipation (MA) = ((total activity 3 h prior to lights-on)/(total activity 6 h prior to lights-on)) 2- (0.5)(ttest p<0.05) for the second Day in DD. **(F)** *060B>UAS-GFP* projections (green) are in close apposition to PDF-labelled sLNv projections (magenta). **(G)** Field of view for Epac recordings. **(H)** FRET ratio of *060B>UAS- Epac1* in response to (1^e-6^M) Pigment dispersing factor (PDF); measured in morning and afternoon (n=4/time of day). **(I-J)** Pre-incubation of *060B>Epac1* with 1^e-6^M PDF dramatically increases the amplitude of 060B responses to 3^e-5^ M DA in the afternoon but not the morning (n= 7-11 neurons/group). **(K)** Quantification of I and J;* p<0.05 modified Bonferroni test **(L)** MB ablation increases the response of *060B >UAS-Epac1* to 3^e-5^ M DA in the PM vs AM; (n=7-17neurons/ condition). **(M )** Gaboxadol restores afternoon responses to DA following HU ablation (n=5-17) neurons/ condition). **(N)** Quantification of L and M; ;* p<0.05 modified Bonferroni test. **(O)** MB ablation increases the response of *051B >UAS-Epac1* to 3^e-5^ M DA in the PM; (n=8 neurons/ condition). **(P)** Gaboxadol restores afternoon responses to DA following HU ablation (n=7 neurons/ condition). **(Q)** Quantification of O and P;* p<0.05 modified Bonferroni test. **(R)** Fas2 staining in *060B/+, UAS- Pdfr^RNAi^/+ and 060B>Pdfr^RNAi^* flies. **(S)** MB ablation disrupted sleep in both parental controls and experimental flies (n=10-16 flies/ condition; *p<0.05 modified Bonferroni test. **(T)** *MB-ablated 060B>Pdfr^RNAi^* flies form an intact STM compared to vehicle fed siblings; MB- ablated parental lines are learning impaired; n>8 flies/condition;* p<0.05 modified Bonferroni test**. (U)** Vehicle-fed *Pdfr^5304^* mutants display learning impairments in the morning n= 7-18 flies/condition ;* p<0.05 modified Bonferroni test**. (V)** Fas2 staining in vehicle and HU fed *Pdfr^5304^* flies. **(W)** Vehicle-fed *Pdfr^5304^* mutants display learning impairments in the morning; HU ablated *Pdfr^5304^* Learn in both the AM and PM, n= 7 flies/condition ;* p<0.05 modified Bonferroni test. Statistics are reported in Supplemental Table 4.

### PDF alters the response properties of γ2α’1-projecting DANs

To determine whether γ2-projecting *060B* DANs could be influenced by the clock, we expressed *UAS-GFP* in γ2- projecting *060B* DANs and co-stained with anti-Pigment dispersing factor (PDF). As seen in Figure 3F-G, PDF dorsal axonal projections can be seen to pass very close to the cell bodies of *060B* neurons. To assess the functional relationship between the LNvs and *060B* neurons, we used live-brain imaging to define cAMP response properties in individual *060B* neurons to bath-applied PDF. Individual traces from single neurons in response to DA or PDF are shown in Supplemental Figure S2 D-E. Interestingly, only 2 of the 8 neurons labelled by the 060B driver responded to PDF, and these responses were larger in the AM (Figure 3H). To determine whether the time-of-day response of *060B* neurons to DA was influenced by PDF, we pre-incubated brains with 1^e-6^ M PDF before examining the response to DA (*104*). In the morning, pre-incubation with PDF had little if any effect on max FRET amplitude following bath-applied DA (Figure 3I). However, in the afternoon, pre-incubation with PDF shifted the response of *060B* neurons to resemble morning responses (Figure 3J-K). These data are consistent with the literature showing that PDF can differentially stagger the timing of activity peaks in diverse neuronal groups to create multiple phasic time points of activity (*105*). Thus, PDF alters the response properties of γ2α’1-projecting DANs.

### PDF plays a role in the time-of-day regulation of STM

Based upon the data presented above, we hypothesized that expressing the *Pigment-dispersing factor receptor* (*UAS-Pdfr^WT^*) in PPL1-γ2α’1-projecting DANs should disrupt STM in the afternoon. Since previous studies have shown that PDF can act on DANs to regulate clock-related behaviors, we began by examining locomotor activity in *060B>Pdfr^WT^* flies and their parental controls (*060B/+* and *UAS-Pdfr^wt^/+*) (*35, 36*). Constitutively expressing *Pdfr^WT^* in *060B* neurons dramatically reduces the percentage of rhythmic flies and disrupts morning anticipation (Supplemental Figure S3 F-K). Importantly, *060B>Pdfr^WT^*flies form an STM in the AM but do not form an STM in the PM; *060B/+* and *UAS-Pdfr^w^/+^t^*, parental controls exhibit an STM in both the AM and PM (Supplemental Figure S3L). Sensory thresholds were not changed in *060B>Pdfr^WT^* flies, indicating changes in learning are not due to changes in the way the animals perceive light or quinine (Supplemental Figure S3 M). These data indicate that PDF plays a role in the time-of-day regulation of STM through its influence on γ2α’1- projecting DANs.

### Knocking down the Pdfr in γ2α’1-projecting DANs prevents social jet lag induced deficits in STM

To better understand the role of PDF in regulating STM, we evaluated flies using a recently developed social jet lag protocol in which flies experience a 3 h light phase delay on Friday followed by a 3 h light advance on Sunday (*80*). This protocol results in persistent desynchrony in PERIOD oscillations in most circadian neurons (*80*). Importantly, social jet lag results in long-lasting impairments in STM after the flies are returned to their typical weekday schedule (*106*). We hypothesize that knocking down the *Pdfr* in γ2α’1-projecting DANs will prevent social jet lag from disrupting STM. As seen in Supplemental Figure S3N, sleep levels in *060B/+>Pdfr^RNAi^/+* and their parental controls, return to their typical weekday schedules after social jet lag as previously reported. Importantly, social jet lag disrupts STM in both *060B/+* and *UAS-Pdfr^RNAi^/+* parental controls but STM is unaffected in social jet lagged *060B/+>Pdfr^RNAi^/+* flies (Supplemental Figure S3O). Social jet lag does not alter photosensitivity or quinine sensitivity consistent with previous reports (Supplemental Figure S3P). Thus, PDF plays a role in the STM deficits induced by social jet lag.

### Altering the timing of activity in γ2α’1-projecting DANs disrupts STM

In order to gain deeper insights into the functional implications of activating γ2α’1-projecting DANs during a less favored time of day, we induced the expression of *UAS-TrpA1* in *060B* neurons and raised the temperature to 31°C for 4 hours every morning or afternoon, continuously for three consecutive days (Supplemental Figure S3, Q-R). Flies were returned to 25°C each day, and STM was examined in the AM on Day 4. Activating *060B/+>UAS-TrpA/+* neurons in the AM did not result in changes to STM compared to *060B/+* or *UAS-TrpA1/+* parental controls (Supplemental Figure 3Q). However, activating *060B/+>UAS-TrpA/+* neurons in the PM disrupted STM compared to *060B/+* and *UAS-TrpA1/+* parental controls (Supplemental Figure S3R). Thus, altering the timing of activity of γ2α’1-projecting DANs can induce deficits in STM.

### MB-ablation disrupts, and sleep restores the time of day γ2α’1-projecting MBONs and DANs are responsive

Given that the activity of γ2α’1-projecting DANs has the potential to modify STM, and that MB ablation can alter the response characteristics of clock neurons (Supplemental Figure S3C), we conducted live-brain imaging to assess the impact of MB ablation on *060B* and *051B* neurons during both the morning and the afternoon. As seen in Figure 3L, MB ablation eliminates the morning/afternoon difference in the response of *060B* neurons to DA. However, the morning/evening response properties of MB-ablated *060B/+>UAS-Epac1-camps* flies is restored after 2 days of Gaboxadol-induced sleep (Figure 3M-N). A similar result was observed for γ2α’1- projecting MBONs (Figure 3O-Q). Thus, sleep can restore baseline responses in γ2α’1-projecting MBONs and DANs to MB-ablated flies.

### Knocking down the Pdfr in γ2α’1 projecting DANs prevents STM impairments in MB-ablated flies

Based upon the data presented above, PDF is, at least in part, a reasonable candidate for mediating STM deficits in MB-ablated flies. Thus, we hypothesized that knocking down the *Pdfr* in γ2α’1-projecting DANs would prevent MB ablation from disrupting memory. To test this hypothesis, we fed HU to *060B/+*, *UAS-Pdfr^RNAi^/+* and *060B/+>UAS-Pdfr^RNAi^/+* larvae and evaluated behavior in adults. As seen in Figure 3R, HU similarly eliminated most Kenyon Cells in each genotype. Moreover, HU-treated flies showed similar reductions in sleep (Figure 3S). Despite expressing *UAS-Pdfr^RNAi^* in only 8 neurons, MB-ablated *060B/+>UAS-Pdfr^RNAi^/+* flies display an intact STM in both the AM and PM (Figure 3T, Supplemental Figure S3, P). Importantly, STM was impaired in both MB-ablated *060B/+* and *UAS-Pdfr^RNAi^/+* parental controls at each time point (Figure 3T, Supplemental Figure S3S). No changes in sensory thresholds were observed, indicating that the changes in STM are not due to changes in sensory thresholds (Supplemental Figure S3, T-U). Thus, STM can be restored in MB-ablated flies by preventing γ2α’1-projecting DANs from receiving a crucial clock signal.

### MB ablation can restore STM to impaired *Pdfr^5304^* mutants

In comparison to *060B/+>UAS-Pdf^WT^/+* flies that have intact MBs and exhibit STM impairments specifically in the PM, sham-treated *060B/+>UAS-Pdfr^RNAi^/+* flies demonstrate normal STM in both the AM and PM, as indicated by the black bars in Figure 3T and Supplemental Figure S3S. Together with recent reports, these data suggest that the *Pdfr* might influence STM via additional MB circuits (*34*). Thus, to better understand the role of *Pdfr* on STM, we examined *Pdfr* mutants (*Pdfr^5304^*) and their *Cs* controls. Consistent with the data for all the control genotypes presented thus far, *Cs* flies form an STM in the AM and PM (Figure 3U). However, STM is impaired in *Pdfr^5304^* mutants in the AM, while their siblings form an STM in the PM (Figure 3U). Although these data highlight the role of *Pdfr* for circuits supporting STM in the AM, it leaves open questions about how MB ablation might alter STM in memory-impaired flies lacking the *Pdfr*. Thus, we examined STM in *Pdfr^5304^* mutants and their *Cs* controls after MB-ablation. As seen in Figure 3V, HU eliminated most Kenyon Cells in both genotypes. Moreover, sleep was severely disrupted in HU-treated *Pdfr^5304^* mutants compared to sham-treated controls (Supplemental Figure S3V). Consistent with previous results, no STM was observed in MB-ablated *Cs* flies (Figure 3W) (*41*). Remarkably, MB-ablated *Pdfr^5304^* mutants display an STM in both the AM and PM (Figure 3W). No changes in sensory thresholds were observed for either genotype (Supplemental Figure S3, W-X). These data reveal a startling observation: MB ablation can restore STM in impaired *Pdfr^5304^* mutants.

### MB ablation and Gaboxadol restore STM to the canonical clock mutant *period* (*per^01^*)

A growing body of evidence suggests that the differential modulation of appetitive and aversive circuits is required for memory acquisition (*16, 95*). Our data indicate that MB ablation alters the timing in the AM-circuit in flies with an intact clock (Figure 3L-Q). Thus, we predict that in flies without a clock the coordination of appetitive and aversive circuits will be governed independently by increased sleep drive (Gaboxadol) or increased wake drive (MB ablation). To test this hypothesis, we first examined STM in *per^01^* mutants in LD and DD. Consistent with previous reports, *per^01^* mutants displayed an STM in LD but were impaired in DD (Supplemental Figure S4A)(*92, 107*). Since *per^01^* mutants have disrupted STM specifically in DD, we evaluated the effects of enhanced sleep or MB ablation in DD. Larvae were fed HU or sham-treated and behavior was evaluated in adults. As seen in Figure 4A, KCs were severely disrupted compared to sham-treated controls. Importantly, MB-alation severely disrupted sleep in *per^01^* mutants whereas Gaboxadol increased sleep in MB- intact *per^01^* flies as expected (Figure 4B). As seen in Figure 4C, we replicated the STM results for intact *per^01^* mutants in LD and DD. Importantly, both increased sleep and MB ablation restored STM to *per^01^* mutants in DD despite producing opposite phenotypes (Figure 4C). No STM was observed when *per^01^* flies were sleep deprived while on Gaboxadol (Supplemental Figure S4B). Given the surprising nature of these results, we examined STM in MB-ablated *per^01^* mutants in LD. As seen in Supplemental Figure S4C, MB-ablation disrupted STM in *per^01^* mutants both in the AM and PM. Neither MB-ablation nor Gaboxadol altered sensory thresholds (Supplemental Figure S4D-E). These data suggest that, in the absence of a circadian clock, sleep drive or MB ablation (perhaps through increasing wake drive) can support the appropriate modulation of appetitive and aversive circuits sufficiently to allow STM in the APS.

**Figure 4:**
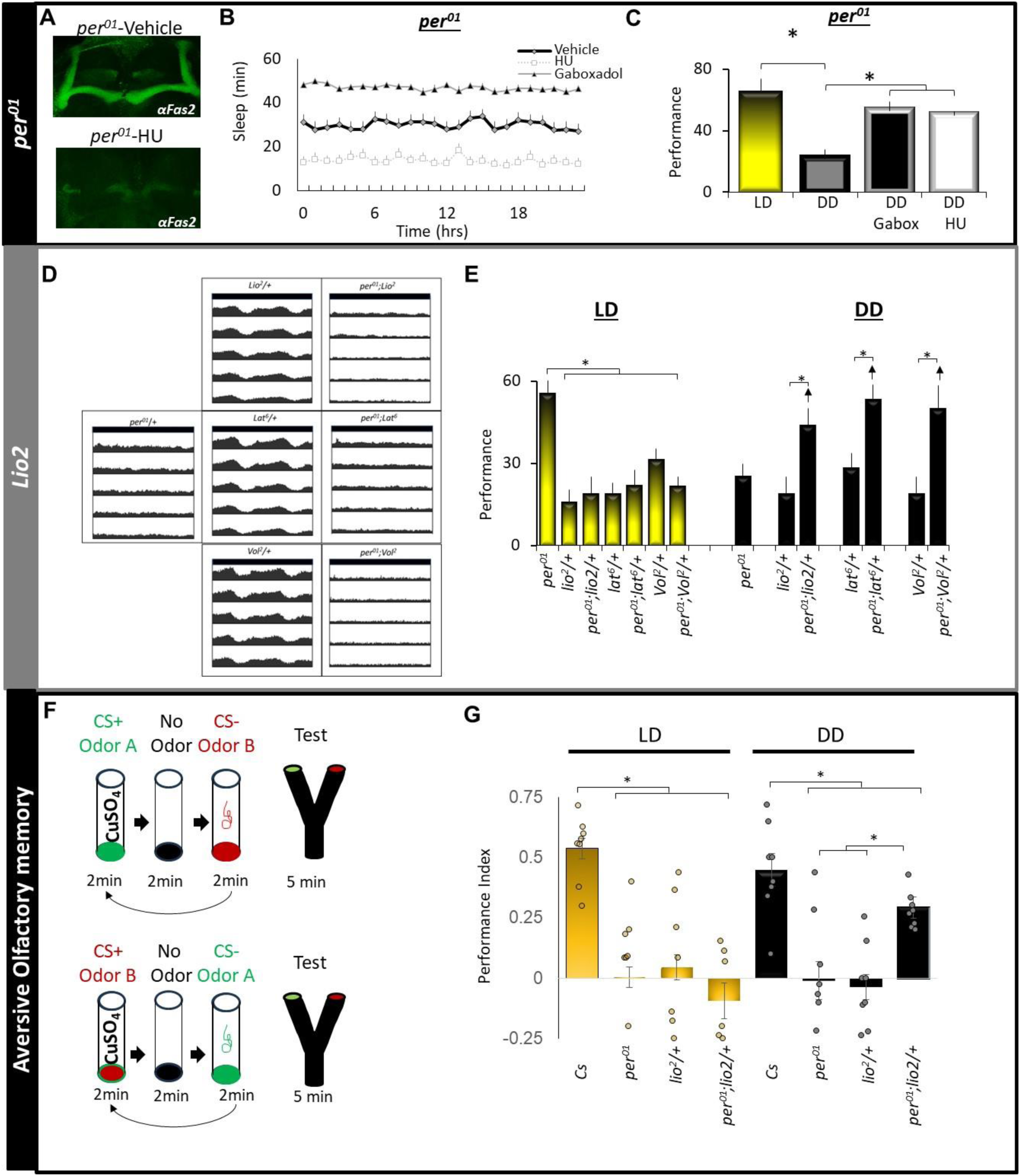
Both HU ablation and sleep restore learning to *per^01^* mutants. **(A)** Fas2 staining in vehicle and HU fed *per^01^* flies **(B)** MB ablation reduces sleep and Gaboxadol increases sleep in *per^01^* flies maintained in n= 30-60 flies/condition. **(C)** Both MB-ablation and Gaboxadol restore learning to *per^01^*flies tested in DD. (n= 8-14 flies/condition; *modified Bonferroni test p>0.05). **(D)** Actograms showing average activity in constant darkness; data is double-plotted. (n=28-32 flies/condition) **(E)** In LD, *per^01^* flies display an STM but no STM is observed in *lio^2^*/+, *per^01^*;*lio^2^*/+, *lat^6^*/+, *per^01^*;*lat^6^*/+, *vol^2^*/+ or *per^01^*;*lat^6^*/+, flies. In DD, no STM is observed in *per^01^* flies or in in *lio^2^*/+, *lat^6^*/+, or *vol^2^*/+ mutants; *per^01^*;*lio^2^*/+, *per^01^*;*lat^6^*/+, and *per^01^*;*lat^6^*/+ flies all display STM (7-10 flies/condition, *modified Bonferroni test p>0.05). **(F)** Y-maze schematic. **(G)** In LD CS flies show immediate aversive olfactory memory while no STM is observed in *per^01^* , *lio^2^*/+, and *per^01^*;*lio^2^*/+. In DD, and STM is observed in only observed in Cs and *per^01^*;*lio^2^*/+ flies (8-18 flies/condition, *modified Bonferroni test p>0.05). Statistics are reported in Supplemental Table 5.

### The *per^01^* mutation restores STM to classical memory mutants only in DD

The circadian modulation of neural circuits in healthy animals is important for maintaining a variety of adaptive behaviors. However, our data suggest that in some circumstances, activity patterns in unhealthy brains (e.g. MB ablation) may place neural circuits into conflict with the timing imposed by the clock. Under such circumstances, removing the putative conflict should allow for STM to proceed normally. To test this hypothesis, we re-examined three classic memory mutants, *lio^2^*, *Origin recognition complex subunit 3* (*lat^6^*) and *scab* (*scb^Vol2^*). As seen in Supplemental Figure S1, and Figure S4 F-G, each of these mutants is memory impaired using the APS thereby replicating results obtained using aversive olfactory conditioning (*77, 79, 108*). Importantly, Gaboxadol-induced sleep successfully restored STM in each case indicating that the STM deficits may be due to changes in neuromodulation rather than deficits in encoding. Since each mutant displayed STM impairments as heterozygotes, we crossed them to *per^01^* females and evaluated them in both LD and DD. As seen from the actograms in Figure 4, B and Supplemental Table 6, no circadian rhythms were observed in *per^01^*; *lio^2^/+*, *per^01^*; *lat^6^/+* or *per^01^*; *scb^Vol2^/+* maintained in DD. When maintained in LD, only *per^01^* flies displayed an STM, while all other genotypes remained impaired (Figure 4E). However, a different pattern emerged for flies maintained in DD. Specifically, no STM was observed when a mutant was tested on its own in DD (Figure 4E). Importantly, while *per^01^*could not restore STM to these memory mutants in LD, *per^01^*; *lio^2^/+*, *per^01^*; *lat^6^/+* or *per^01^*; *scb^Vol2^/+* all restored STM in DD (Figure 4E). A similar picture emerged when using the Y-maze to evaluate immediate aversive olfactory memory in *per^01^*, *lio^2^*/+ flies (Figure 4F,G). Together these data highlight the importance of neuromodulation for restoring STM in flies with mutations in genes that play important roles in memory formation.

## Discussion

We report that sleep can restore memory to MB-ablated flies and that MB-ablation does not produce memory impairments in clock mutants maintained in constant conditions. Importantly, we demonstrate that neurons that are predominantly involved in aversive learning (e.g. γ2-projecting MBONs/DANs) support memory in the morning, while neurons that are predominantly involved in appetitive learning (e.g. γ5 projecting MBONs/DANs) support memory in the afternoon (*16*). During MB-ablation the responses of morning active γ2 projecting MBONs/DANs persist into the afternoon, potentially creating a conflict between aversive and appetitive circuits (*12*). We identify *Pdfr* as a potential mediator for the changes in γ2 projecting DANs during MB-ablation. Importantly, we find that selectively knocking down the *Pdfr* in these eight DANs effectively restores memory to MB-ablated flies. These data indicate that MB-ablation disrupts memory, in part, through its impact on the clock.

Although the relationship between the MBs and clock circuitry is well established (*98, 99, 109*), we did not expect the clock to be involved in memory deficits induced by MB-ablation. Thus, we conducted three independent experiments to evaluate the role of the clock on memory as assessed using the APS. First, we demonstrated that constitutively expressing *UAS-Pdfr^WT^* in γ2 projecting DANs dramatically reduces the percentage of rhythmic flies, alters morning anticipation, and disrupts STM in the afternoon. These data highlight the interaction between clock and MB circuitry (*35*). Secondly, we knocked down the *Pdfr* in γ2 projecting DANs and exposed the flies to social jet lag. Social jet lag results in persistent desynchrony in PERIOD oscillations in most circadian neurons and impairs STM (*80*). Knocking down the *Pdfr* in γ2 projecting DANs prevented social jet lag from inducing STM deficits, further linking clock dysfunction with MB circuitry. Thirdly, we selectively increased the activity of γ2 projecting DANs in the morning or in the afternoon. Memory impairment was only observed when the activity of γ2 projecting DANs was displaced into the afternoon. These data highlight the role that clock circuits play in modulating MB-circuits and thus STM in the APS.

To map the circuits underlying STM in the APS, we expressed wild-type *rutabaga* in γ lobe-projecting KCs, MBONs and DANs in flie’s mutant for *rutabaga* in all other neurons (*89, 90*) Our data yielded two surprising results. First, STM could be rescued when UAS-*rut^WT^* was expressed at any level of the circuit hierarchy. For example, STM was restored when *rutabaga* was restored in MBONs even though upstream KCs and DANs were mutant for *rutabaga;* neither *rut^2080^* mutants nor *rut^2080^;GAL4/+* controls learn in the morning or afternoon. Similar results were observed when rescuing *rutabaga* in light and quinine sensing neurons even though all downstream neurons are mutant for *rutabaga*. These findings reveal that restoring a single gene, *rutabaga*, in specific neurons can compensate for genetic deficits across the broader circuit. Understanding the molecular mechanisms behind this non-cell autonomous rescue may provide insight in to treating cognitive and neurological disorders and is the subject of active investigation.

Second, the switch from learning to not learning was abrupt. For manipulations to the morning circuits, flies would robustly form an STM from 8am until ∼12pm and then suddenly be unable to form an STM. The converse was true for manipulation to the afternoon circuit; flies would be unable to form an STM between 8am to ∼12pm and then suddenly acquire the ability to form an STM. It is unlikely that the constitutive or ectopic expression of *rutabaga* was the cause, as complementary results were observed when *UAS-Dop1R1^RNAi^*was used to disrupt these circuits. The reason why the clock prioritizes aversive circuits for STM in the morning and appetitive circuits in the afternoon remains unclear. However, one hypothesis is that the clock has evolved to align specific types of learning valence with ethologically predictable situations, such as avoiding predators in the morning and foraging in the afternoon, for example. Under this scenario, the clock could indirectly regulate memory circuits through its effects on metabolic centers. However, the persistence of time-of-day segregation of aversive and appetitive circuits during starvation, does not support this hypothesis as metabolic demands are uniformly high throughout.

An alternative hypothesis is that the clock is playing a direct and active role in determining when a circuit can be used for STM. A growing body of evidence suggests that normal waking experience progressively disrupts neural circuits, and that sleep can restore these deficits (*110, 111*). Animals may not have the opportunity to sleep during their primary wake-period resulting in the functional degradation of a circuit and reduced fitness. We hypothesize that the circadian clock plays a critical role in preventing circuit failure by strategically regulating activity: it turns certain circuits off to avoid overuse and degradation, and then activates “rested” circuits at specific times, ensuring optimal neural function during waking hours. This hypothesis will be tested in future studies.

Finally, we explored the relationship between the clock, sleep, and MB-circuits by examining STM in *per^01^* mutants during MB ablation, sleep enhancement and in the context of classical memory mutants. Mutants for *per^01^* learn in LD but are impaired DD. Enhancing sleep restores STM to MB-intact *per^01^*mutants in DD. Conversely, STM is disrupted in short-sleeping, MB-ablated *per^01^* mutants in LD but is restored in DD. Similarly to MB-ablation, STM is restored in *per^01^; lio^2^/+*, *per^01^*; *lat^6^/+*, and *per^01^*;scb^Vol2^*/+* flies but only in DD. Together, our data suggest that the temporal organization of appetitive and aversive circuits in brain damaged flies, or in some memory mutants, can be compensated for by increased sleep or by eliminating conflicting signals emanating from the clock. These observations are consistent with previous reports showing that ablating the suprachiasmatic nucleus in arrhythmic Siberian hamsters and in a mouse model for Downs syndrome (Ts65Dn) restores memory (*112, 113*). Together with our observations in flies, these data suggest that ability of clock circuits to amplify cognitive impairments in the context of circuit dysfunction may be evolutionarily conserved.

In conclusion, our study not only highlights the intricate interplay between circadian rhythms, sleep, and memory in fruit flies, it also emphasizes the utility of using enhanced sleep as tool to evaluate sleep’s function. By observing how sleep restored memory in brain-damaged flies, we uncovered unexpected molecular pathways and neuromodulatory mechanisms. These insights allowed us to manipulate molecular pathways in a similar manner as enhanced sleep, but without directly inducing sleep. Such an approach provides a pathway towards harnessing the therapeutic potential of sleep for treating cognitive and neurological disorders.

## Methods

### EXPERIMENTAL PROCEDURES

#### Flies

Flies were cultured at 25°C with 50-60% relative humidity and kept on a diet of yeast, dark corn syrup and agar under a 12-hour light:12-hour dark cycle. *104y-GAL4*, flies were obtained from M. Heisenberg (Rudolf Virchow Center). *UAS-NaChBac*(*71*) flies were obtained from A. Sehgal (University of Pennsylvania). *rut^2080^*and *rut^2080^; UAS-rut^WT^* were obtained from Troy Zars (University of Missouri). *CK2β^mbu^* obtained from Thomas Raabe (University of Wurzburg) *Pdfr*-null mutant (*Pdfr^5304^*)(*114*) (RRID:BDSC_33068) ; *UAS-Pdfr^wt^ ; w; UAS-Epac1camps50A;* UAS-*Pdfr^RNAi^*and *PDF-GAL4* were obtained from Paul Taghert (Washington University in St. Louis). MB334C(*115*) was obtained from Jerry Rubin (Janelia Farms Research Campus). GRASP reagents – *UAS CD4:SpGFP_1-10_*, LexAop *CD4:SpGFP_11_*, *UAS brp:mcherry* (3^rd^ chromosome insert), *lexaop brp:mcherry* (3^rd^ chromosome insert) were gifts of C-H. Lee (Academia Sinica, Taiwan)(*116–118*). LexAop P2X2(*119*) was a gift of O. Shafer (U. Michigan). The following stocks were obtained from the Bloomington Stock center: w[1118]; Df(2R)lio[2](*120, 121*), pigeon[2] drl[2]; w1118; P(*85*)pigeonP1, drlP1/CyO, P(sevRas1.V12}FK1; w1118; P(UAS-RA.cs2}39 (RRID:BDSC_38623)(*84*); MB247-GAL4 ; 201y-GAL4; *Dop1R1RNAi^HMC05200^* (RRID:BDSC_62193)*; Dop1R1^RNAiHM04077^* (RRID:BDSC_31765)*; Dop1R2 ^RNAiHMC06293^* (RRID:BDSC_65997)*; DopEcR^RNAiJF03415^* (RRID:BDSC_31981) *; D2R^RNAiHMC02988^* (RRID:BDSC_50621), *AstaR1^RNAi^* (RRID:BDSC_27280) (RRID:BDSC_50621); MB131B (RRID:BDSC_68265); MB607B (RRID:BDSC_68256); MB112C (RRID:BDSC_68263); MB438B (RRID:BDSC_68326); MB051B (RRID:BDSC_68275); 060B, (RRID:BDSC_68279); MB296B (RRID:BDSC_68308); MB083C (RRID:BDSC_68287); MB025B (RRID:BDSC_68299); MB441B(RRID:BDSC_68251); MB434B (BDSC_68325); MB312C, (RRID:BDSC_68252); MB210B (RRID:BDSC_68272); MB011B (RRID:BDSC_68294); MB313C (RRID:BDSC_68315); MB334C, GR66a-GAL4(RRID:BDSC_57670)(*16, 38*) .

#### Sleep

Sleep was assessed as previously described (*58*). Briefly, flies were placed into individual 65 mm tubes and all activity was continuously measured through the Trikinetics Drosophila Activity Monitoring System (www.Trikinetics.com, Waltham, Ma). Locomotor activity was measured in 1-minute bins and sleep was defined as periods of quiescence lasting at least 5 minutes.

#### Pharmacology

Gaboxadol was administered at a dosage s 0.1 mg/mL in standard fly food. Flies were maintained on the drug for the durations described in the text during which time sleep was monitored. Flies were removed from Gaboxadol one hour prior to being tested for short-term memory and one hour prior to being trained for courtship conditioning.

#### Short-term memory

Short-term memory (STM) was assessed by Aversive Phototaxic Suppression (APS) as previously described (*41, 70*). The experimenters were blinded to condition. In the APS, flies are individually placed in a T-maze and allowed to choose between a lighted and darkened chamber over 16 trials. Flies that do not display phototaxis during the first block of 4 trials are excluded from further analysis (*70, 122*). During 16 trials, flies learn to avoid the lighted chamber that is paired with an aversive stimulus (quinine/humidity). The performance index is calculated as the percentage of times the fly chooses the dark vial during the last 4 trials of the 16 trial test. In the absence of quinine, where no learning is possible, it is common to observe flies choosing the dark vial once during the last 4 trials in Block 4 (*70*). In contrast, flies never choose the dark vial 2 or more times during block 4 in the absence of quinine (*70*). Thus, STM is defined as two or more photonegative choices in Block 4. Power analysis using G*Power calculates a Cohen’s d of 1.8 and indicates that 6-8 flies/group are needed to obtain statistical differences (*70*). Within a group of flies that from an STM, ∼6/8 flies will choose the dark vial 2 times in block 4 while ∼2/8 will choose the dark vial 1 time in block 4; the reverse pattern is observed for cohorts of flies that do not form an STM (*4, 69, 70*). While small variations of this pattern are observed, the distribution of performance scores remains essentially quantal. These observations highlight the distinct behavioral patterns between these two groups. Based on these observations, plotting individual data points on top of the group mean would not provide any additional meaningful information. Instead, it would result in cluttered and visually complex figures, potentially obfuscating the key patterns and trends that we aim to communicate. To ensure transparency and accessibility of the data, we include the scores for individual flies in a supplemental File 1.

#### Courtship Conditioning

Training for 4–8 day old males was based on previously described methods (*4, 58*). The males were exposed to mated females in a training protocol consisting of three one-hour training sessions, each separated by one hour. Long-term memory was tested forty-eight hours after the beginning of training, when trained and naive males were exposed to virgin females for a 10-minute testing period (n=16-30 flies/condition). The Courtship Index (CI) is defined as the percent of time that each subject fly spends in courtship behavior during the 10- minute testing period. The CIs were subjected to an arcsine square root transformation to approximate normal distribution as described in (*123*). Data are presented as a Performance Index (PI), where PI=((CI_average-naive_-CI_Trained_)/CI_average-naive_) X100); PIs were evaluated using the Kruskal-Wallis test. The experimenters were blinded to condition. For the Space training experiments with Gaboxadol-induced sleep, naïve males were fed 0.1G for 48 h prior to training. Gaboxadol-fed flies were removed from Gaboxadol 1 h prior to and during training and then returned to 0.1G for 24 h post-training as previously described (*4*). Vehicle-fed flies were maintained on vehicle throughout the protocol.

#### Photosensitivity

Photosensitivity was evaluated as previously described(*70*). Briefly, flies were put in the T-maze over 10 trials in the absence of filter paper. The lightened and darkened chambers appeared equally on both the left and right. The photosensitivity index (PI) is the average of the scores obtained for 5-6 flies ± s.e.m.

#### Quinine sensitivity

Quinine sensitivity index (QSI) was evaluated as previously described (*41, 70*). Briefly, flies were individually placed at the bottom of a 14 cm transparent cylindrical tube which was uniformly lighted and maintained horizontal after the introduction of the animal. Each half of the apparatus contained separate pieces of filter paper which could be wetted with quinine or kept dry. The QSI was determined by calculating the time in seconds that the fly spent on the dry side of the tube when the other side had been wetted with quinine, during a 5 min period.

#### Aversive Olfactory Conditioning (Y-maze)

Flies were trained with either Odor A (MCH) or Odor B (OCT) as the conditioned stimulus (CS+). During training, flies experienced 80 mM CuSO4 (US) with the CS+. Immediate memory was evaluated by measuring the preference for CS+ and the alternative CS-. The Performance Index (PI) = (CS+ choices - CS- choices)/total flies). Scores from the fly groups trained with OCT or MCH as CS+ and were averaged to derive a data point.

#### Imaging

Methods generally followed those of (*96*)Klose et al., (2016). Flies were removed from DAM monitors and glass tubes were placed on ice for approximately 5 minutes. 3-4 flies were pinned onto a sylgaard dissection dish, and were dissected in cold calcium-free HL3. Dissected brains were transferred onto a poly-lysine treated dish (35 3 10 mm Falcon polystyrene) containing 3 ml of 1.5mM calcium HL3. Two brains were assayed concurrently, typically a mutant line and its genetic controls. Image capture and x,y,z stage movements were controlled using SLIDEBOOK 5.0 (Intelligent Imaging Innovations), which controlled a Prior H105Plan Power Stage through a Prior ProScanII. Multiple YFP/CFP ratio measurements were recorded in sequence from each hemi-segment of each brain in the dish. Following baseline measurements, 1 ml of saline containing various concentrations of either PDF, DA (Sigma-Aldrich) was added to the bath (dilution factor of 1/4). Maximum amplitude values were used to perform ANOVA analyses followed by post hoc Tukey tests. For P2X2 experiments, Gcamp fluorescence images were acquired at 1Hz. Following 1min of baseline measurements, 0.4ml of 5mM ATP was perfused onto the dish to activate the P2X2 receptor.

#### Mushroom Body Ablation

0–1 h old larvae were fed yeast paste (sham-controls) or yeast paste containing 100 mg/mL Hydroxy Urea (HU, Sigma Aldrich, St Louis, MO) for 4 h via standard protocol (*41, 67*). The efficiency of the ablation procedure was evaluated with standard whole mount immunohistochemistry for fasciclin2 (1D4, mouse anti-FASII, 1:75, Hybridoma Bank, University of Iowa) and Alexa 488 anti-mouse IgG (1:1000, Molecular Probes). Confocal stacks were acquired with a 1µm slice thickness using an Olympus FV500 laser scanning confocal microscope and processed using ImageJ.

#### Locomotor Rhythms

Flies were individually placed into 5×65mm tubes with regular food and placed into constant conditions for 6 days. locomotor activity was continuously recorded in 30min bins using the Trikinetics system. Actogram J was used to generate actograms and period measurements(*124*). Flies were scored as rhythmic using the criteria defined by (*125*) by scorers blind to condition. Morning anticipation (MA) = ((total activity 3 h prior to lights-on)/(total activity 6 h prior to lights-on)). Evening anticipation (EA) = ((total activity 3 h prior to lights-off)/(total activity 6 h prior to lights-off)).

#### Social Enrichment

All flies were collected upon eclosion and maintained in same-sex vials containing 30 flies. Flies were divided into a socially isolated group, which were individually housed in 65-mm glass tubes, and a socially enriched group, consisting of 35-45 female flies housed in a single vial. After 5 days of social enrichment/isolation, flies were placed into clean 65 mm glass tubes and sleep was recorded for 3 days as described above. To calculate the effect of social enrichment on subsequent sleep, we first calculate the population mean value for daytime sleep in the isolated group, averaged over 3 days, and then subtracted this isolated group mean from the daytime sleep observed for each individual socially-enriched sibling. The difference is referred to as ΔSleep.

#### Statistics

All comparisons were performed using a Student’s t-test or ANOVA, followed by subsequent planned comparisons employing a modified Bonferroni test, unless specified otherwise. It is important to note that a significant omnibus F-test is not a prerequisite for conducting planned comparisons, as previously established (*126*). Our experimental design is based on the expectation of targeted effects confined to a single group or condition, leaving the remaining groups unaffected (*4*). Consequently, the number of planned comparisons never exceed degrees of freedom (df) df-1. For the circuit mapping experiments shown in Figure 2 and Figure S2, each GAL4 line was crossed and assessed separately and thus constitute independent experiments. All statistically different groups are defined as *p < 0.05.

## Figure Legends

**Associated with Figure 1: Supplemental Figure S1:**
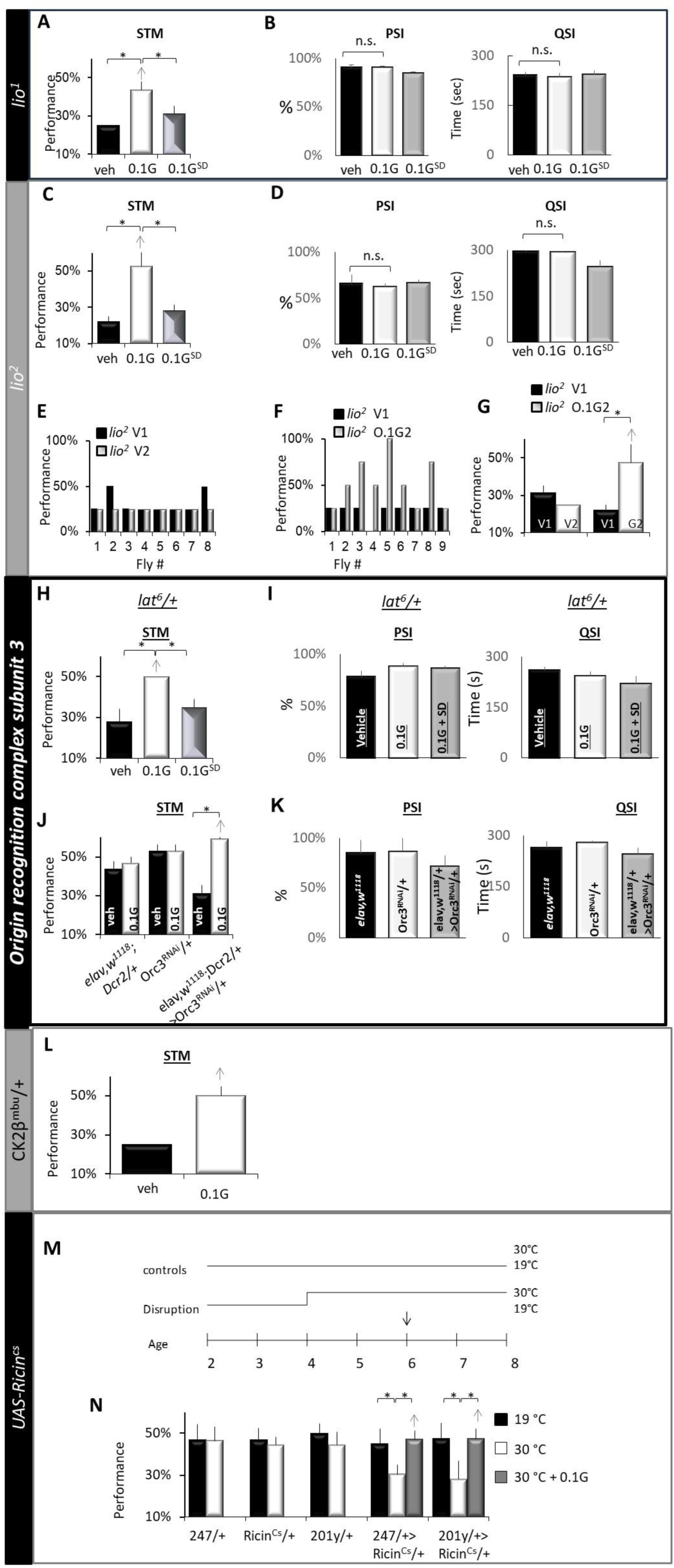
Gaboxadol restores STM to MB disrupted flies. **(A)** *lio^1^*mutants exhibit deficits in STM (veh) which are reversed by Gaboxadol (0.1G); *lio^1^* mutants sleep deprived while on Gaboxadol (0.1G^SD^) did not exhibit STM, *p<0.05 modified Bonferroni test, n=>8/group (see Table 1 for statistics). **(B)** Gaboxadol did not substantially alter PSI, n.s. p>0.05 modified Bonferroni test. **(C)** *lio^2^* mutants exhibit STM deficits (veh) which are rescued by Gaboxadol (0.1G); *lio^2^* mutants sleep deprived while on Gaboxadol (0.1G^SD^) did not exhibit STM, *p<0.05 modified Bonferroni test (n=>8/group). **(D)**. Gaboxadol did not alter PSI One way or QSI. **(E)** Individual *lio^2^* flies maintained on vehicle are deficient in STM when tested on two consecutive trials(V1 and V2). **(F)** The STM deficits in *lio^2^*individuals were reversed following 2 days of Gaboxadol-induced sleep (0.1G2). **(G)** Mean performance scores ± SEM for *lio^2^* flies maintained on vehicle (V1, V2) or switched from vehicle (V1) to Gaboxadol for 2-days (G2); paired t-test, *p<0.05. **(H)** Mutants for *Origin recognition complex subunit 3 (Orc3, lat6)* exhibit STM impairments which are reversed by Gaboxadol induced sleep; no STM is observed when flies fed Gaboxadol are prevented from sleeping; *p>0.05 modified Bonferroni test) **(I)** Gaboxadol did not alter PSI or QSI. **(J)** Knocking down Orc3 pan neuronally (elav>Orc3^RNAi^) disrupted STM compared to parental controls (*elav/+* and *Orc3^RNAi^/+*); Gaboxadol induced sleep restored STM to *elav>Orc3^RNAi^* flies, n= 8 flies/condition; *p>0.05 modified Bonferroni test. **(K)** PSI and Qsi did not differ between *elav>Orc3^RNAi^* and parental controls. **(L)** STM impairments in CK2β^mbu^/+ mutants are reversed by Gaboxadol induced sleep; ttest n=7/8 flies/condition. **(M)** Schematic of experiment. Flies were either maintained at 19°C (black) or placed at 30°C (white) for 4 days. On Day 6, flies expressing the temperature sensitive protein inhibitor *UAS-Ricin^cs^*either throughout the mushroom Body (*247-GAL4*) or in the gamma lobes (*201y-GAL4*) were placed on THIP for 2 days (arrow). **(N)** Gaboxadol-induced sleep restores STM to *247>UAS-Ricin^Cs^*, (n= 10-17 flies/Condition) and *201y>UAS-Ricin^Cs^* (n= 8-9 flies/Condition); Modified Bonferroni test *p < 0.05. Gaboxadol was not administered to parental controls because they did not express STM deficits at either 19°C or 30°C). Statistics are reported in Supplemental Table 2.

**Associated with Figure 2 Supplemental Figure S2.**
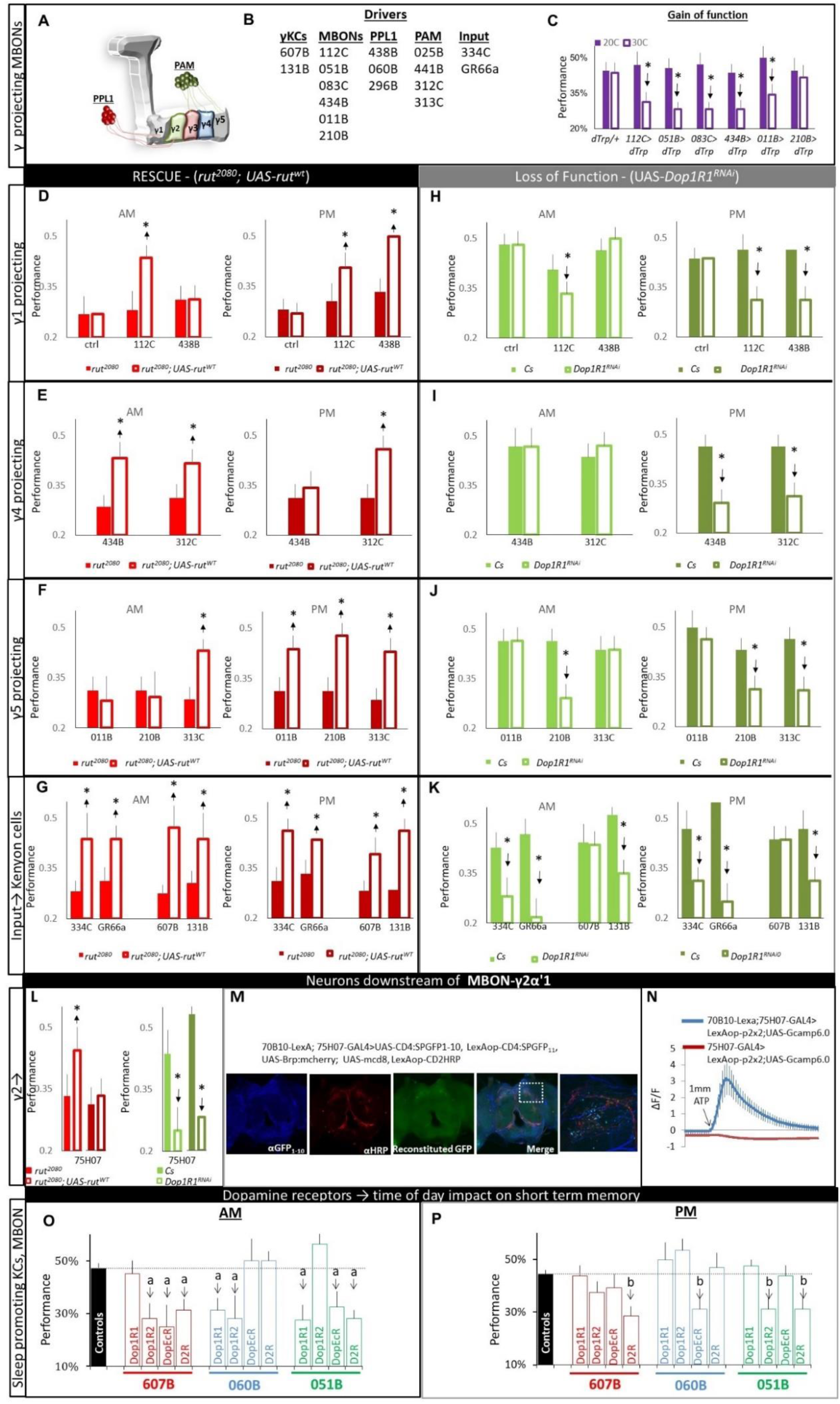
Time of day regulation of STM. **(A)** Schematic of γ lobe compartments and their DANs. **(B)** List of split-GAL4 and GAL4 drivers. **(C)** STM was disrupted in γ lobe projecting MBON Split-GAL4>UAS-TrpA1when the temperature was raised to 30°C overnight; *p<0.05, modified Bonferroni test, (n>8 flies/group) (See Table 2 for statistics). **(D-G, left)** STM was evaluated separately in either the morning (light red) or in the afternoon (dark red). Statistical comparison were made using an ANOVA followed by a modified Bonferroni test to identify differences between experimental (*rut^2080^; split-GAL4>UAS-rut^WT^*) and control flies (*rut^2080^* /+; Split-GAL4/+) for a given time of day (^a^light red bars vs light red bars or ^b^dark red bars vs dark red bars. **(H-K, right)** Short-term memory was also evaluated in flies expressing *UAS-Dop1R1^RNAi^* in individual circuits: *splitGAL4/+>UAS-Dop1R1^RNAi^*/+ vs. parental controls (*splitGAL4/+* and *UAS-Dop1R1^RNAi^*/+ in the morning (light green) or afternoon (dark green), modified Bonferroni test. (n>8 flies/group); ^a, b^ p<0.05) **(L)** STM in the AM (light, color) and PM (Dark color) in *rut^2080^; 75H07- GAL4>UAS-rut^WT^* and *75H07-GAL4>Dop1R1^RNAi^*; modified Bonferroni test. (n>8 flies/group); ^a, b^ p<0.05). **(M)** GFP Reconstitution Across Synaptic Partners, (GRASP), signal showing that *70B10-LexA* neurons which include (*051B* neurons) make contact with *75H07* expressing neurons. **(N)** P2X2 mediated stimulation of *70B10-LexA* neurons by bath application of ATP, increases Gcamp fluorescence in *75H07-GAL4* cell bodies. Control animals lacking the *70B10-LexA* transgene did not exhibit this response (n = 6 cells/condition). **(O)** STM was evaluated in the AM after expressing RNAi for *Dopamine 1-like receptor 2*, *UAS-Dop1R2^RNAi^*, *Dopamine 1-like receptor 2, UAS-DopEcR^RNAi^* and *Dopamine 2-like receptor*, *UAS-D2R^RNAi^* using *607B Split-GAL4*, *060B-Split-GAL4* and *051B- Split-GAL4*. Parental lines were not statistically different and were thus combined; *Dop1R1^RNAi^* data from figure 2 are re-plotted to facilitate comparisons; (^a^p<0.05 modified Bonferroni test vs controls) **(P)** STM was also evaluated in the PM; (^b^p<0.05 modified Bonferroni test vs controls). Statistics are reported in Supplemental Table 3.

**Associated with Figure 3: Figure S3:**
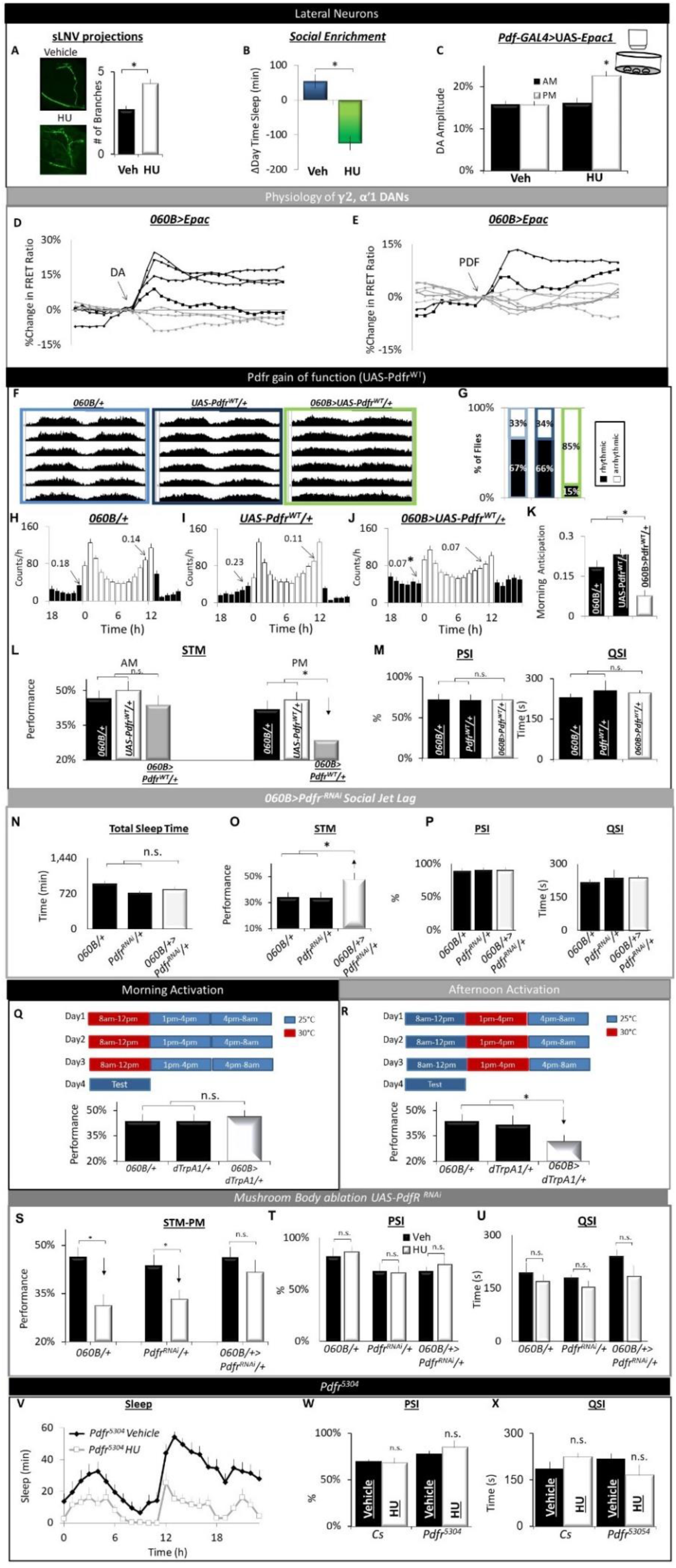
γ2α’1 projecting 060B DANs alter clock-regulated behavior. **(A)** MB-ablated Cs flies and controls were stained for αPDF antibody and the complexity of their branches was quantified as previously described (n=12 hemispheres); ttest, p<0.05. **(B)** Δ Daytime sleep reveals that social enrichment has no impact on MB-ablated Cs flies (green) compared to sham-treated siblings (blue (p= 2.7^E10-5^, Student *t*-test, n = 19-32/group). **(C)** HU ablation increases cAMP in Pdf neurons in response to 3^e-5^ M DA in the afternoon; n= 8-13 neurons/condition,*p>0.05 modified Bonferroni test). **(D)** Individual traces of 060B neurons are not homogeneous in their response to 3^e-5^ M DA; only neurons which exhibit a positive slope considered to respond and are then selected for quantification in the main figures. **(E)** Individual traces of 060B neurons to ^10-6^ PDF. On average only two 060B neurons respond. **(F)** Actograms showing average activity in constant darkness for *060B>Pdfr^WT^* and their parental controls, 060B/+ and UAS-Pdfr^wt^; data is double-plotted. (n=28-32 flies/condition) **(I)** % of rhythmic flies chi-square =19.5; p <0.001. **G-K)** Average hourly normalized activity counts for *060B/+*, *UAS-Pdfr^wt^/+* and 060B/+>UAS-Pdfr^wt^/+ maintained on a Light Dark (LD) schedule (n=29-31 flies/genotype). **(K)** Quantification of Morning anticipation, *p>0.05 modified Bonferroni test. **(L)** *060B>Pdfr^WT^*flies are learning impaired in the PM but not in the AM; *p>0.05 modified Bonferroni test). **(M)** *060B>Pdfr^WT^*does not change PSI or QSI, *p<0.05 modified Bonferroni test. **(N)** Social Jet lag does not alter Total sleep time; n= 10-13 flies/condition * p<0.05 modifid Bonferroni test. **(O)** Social Jet lag disrupts STM in *060B/+* and *UAS- Pdfr^RNAi^/+* but does not disrupt STM in *060B/+>UAS-Pdfr^RNAi^/+* flies; n= 11-12 flies/condition * p<0.05 modified Bonferroni test. **(P)** Social jet lag did not alter PSI; n= 6 flies/condition, or QSI; n= 5 flies/condition. **(Q)** Schematic for 4 h of 060B activation in the AM. Activating 060B neurons for 4 hours in the AM for 3 days, does not disrupt STM the following AM compared to parental controls (ANOVA F_[2,22]_ = 0.14 ; p=0.86; n=7- 8flies/condition; *p>0.05 modified Bonferroni test. **(R)** Schematic for Activating 060B neurons for 4 hours in the PM for 3 days, does not disrupt STM the following AM compared to parental controls (ANOVA F_[2,24]_ = 2.68 ; p=0.09; n=6-11 flies/condition; *p<0.05 modified Bonferroni test). **(S)** *MB-ablated 060B>Pdfr^RNAi^*flies have an STM compared to vehicle fed siblings; MB-ablated *060B/+*, *UAS-Pdfr^RNAi^/+* parental lines are STM impaired; (n>8 flies/condition;* p<0.05 modified Bonferroni test)**. (T,U)** Neither PSI, nor QSI **(V)** HU ablation disrupted sleep in *Pdfr^5204^* compared to sham treated controls (n=12-15 flies/ condition). **(W,X)** No significant differences were observed for PSI and QSI in vehicle or HU fed *Cs or Pdfr^han^* flies (*p<0.05 modified Bonferroni test). Statistics are reported in Supplemental Table 4.

**Associated with Figure 4: Supplemental Figure S4:**
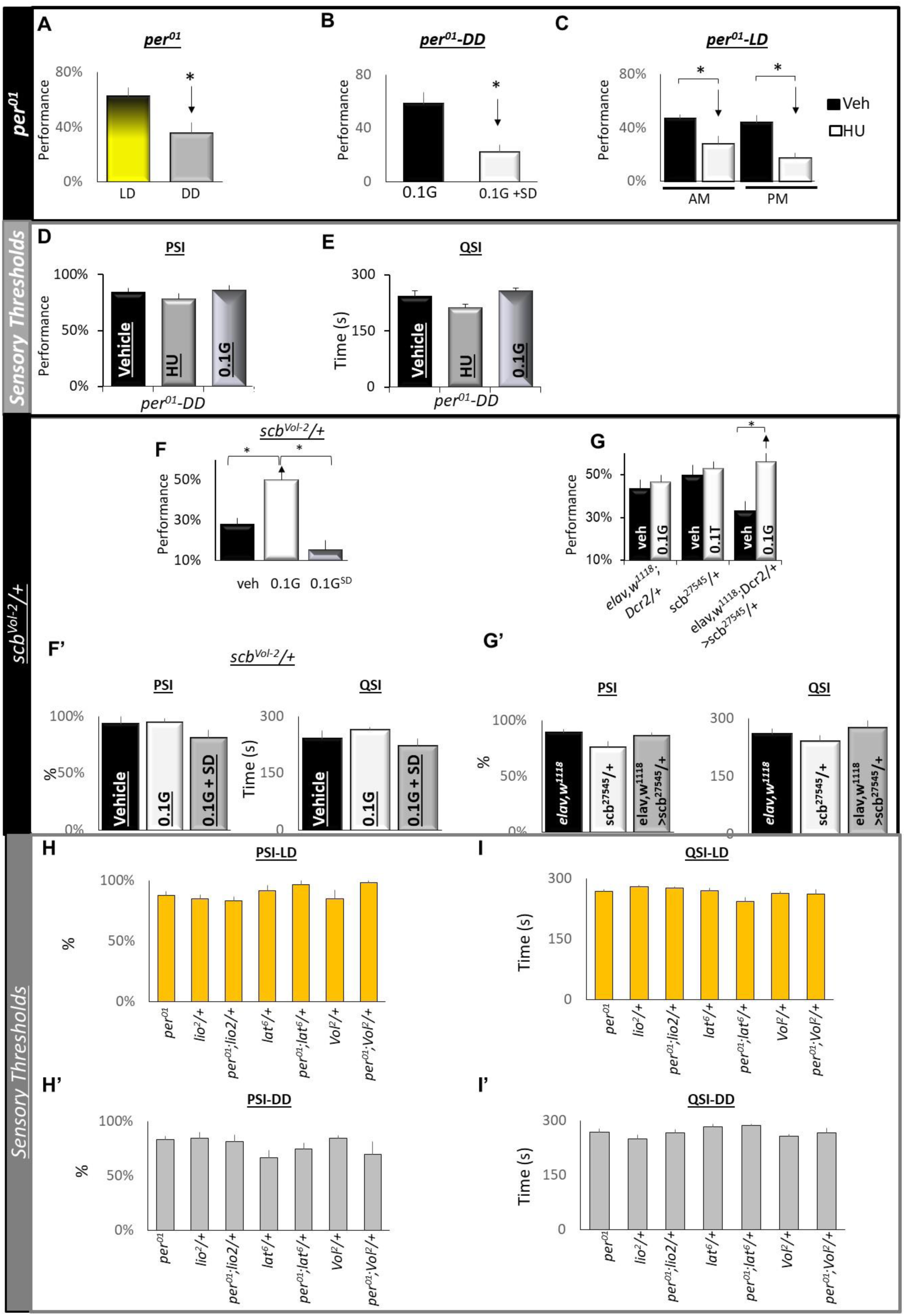
**(A)** *per^01^* flies display an STM in LD but are impaired in DD. (ttest p=0.009, n = 7-10 flies/condition)**. (B)** STM is restored in *per^01^* flies maintained on 0.1G for 2 days in DD; no STM is observed in sleep deprived siblings (ttest p=0.03, n = 3-8 flies/condition. **(C)** HU ablations does not restore STM to in *per^01^* flies maintained in LD in either the AM or PM (n = 8-12 flies/ condition; *p<0.05, modified Bonferroni). **(D,E)** Neither PSI nor QSI differed between either 0.1G fed or HU fed flies (n = 5-6 flies/ condition; *p<0.05, modified Bonferroni). **(F)** Gaboxadol restores STM to *scb^Vol-2^/+* mutants; no STM is observed when flies fed Gaboxadol are prevented from sleeping (n= 8-9 flies/condition; *p>0.05 modified Bonferroni test). **(F’)** Neither PSI nor QSI are altered *scb^Vol-2^/+* mutants. (n = 5-6 flies/ condition; *p<0.05, modified Bonferroni). **(G)** Two days of Gaboxadol-induced sleep restores STM to *elav,w^1118^;Dcr2/+ >scb^27545^/+* compared to Vehicle fed siblings; both parental controls display STM (n= 7-9 flies/condition; *p<0.05, modified Bonferroni). **(G’)** Neither PSI nor QSI is disrupted in *elav,w^1118^;Dcr2/+ >scb^27545^/+* compared to parental controls (n= 5-6 flies/condition; *p<0.05, modified Bonferroni). **(H, H’)** PSI in LD and DD, (6-9 flies/condition, *modified Bonferroni test vs parental control (e.g. *per^01^*;*lio^2^*/+ vs *lio^2^*/+ and *per^01^*, p>0.05). **(I, I’)** QSI in LD and DD, (6-9 flies/condition, *modified Bonferroni test vs parental control (e.g. *per^01^*;*lio^2^*/+ vs *lio^2^*/+ and *per^01^*, p>0.05). Statistics are reported in Supplemental Table 5.

---

**Table S1.**
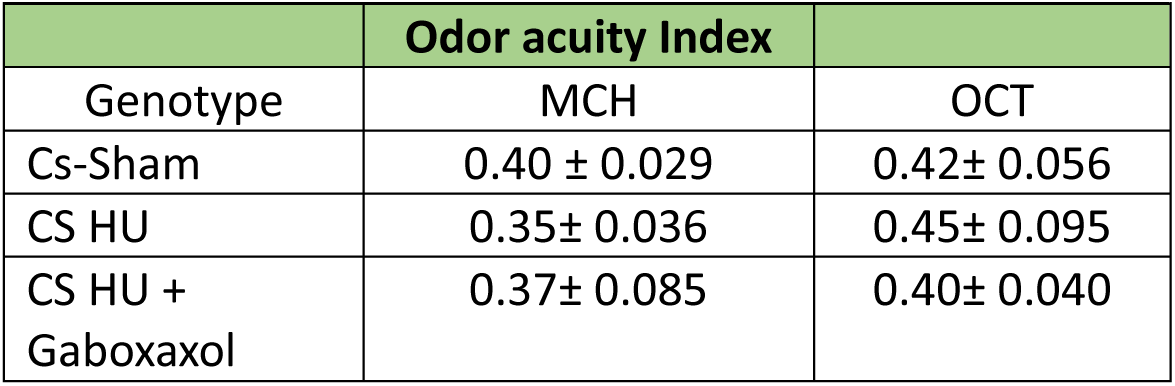

| Table S1 |  |  |
| --- | --- | --- |
|  | Odor acuity Index |  |
| Genotype | MCH | OCT |
| Cs-Sham | 0.40 ± 0.029 | 0.42± 0.056 |
| CS HU | 0.35± 0.036 | 0.45± 0.095 |
| CS HU + Gaboxaxol | 0.37± 0.085 | 0.40± 0.040 |

**Table S2.**

| Table S2 |  |  |  |
| --- | --- | --- | --- |
|  | Statistics for Figure 1 |  |  |
| Figure 1A<br>Cs-Sleep | 2(Veh, HU) X 24(Time) ANOVA $F_{[42,1610]} = 7.47$; $p = 9.9 \times 10^{-16}$ | n=18-26/group | |
| Figure 1B<br>Cs-STM | One way ANOVA for condition (Sham, HU, HU<sup>0.1G</sup>) $F_{[2,22]} = 10.8$; $p = 0.006$ | n=7-8/group | *$p < 0.05$<br>modified Bonferroni test |
| Figure 1B'<br>Cs-PSI | One way ANOVA for condition (Sham, HU, HU<sup>0.1G</sup>) $F_{[2,20]} = 0.95$; $p = 0.4$, | n=5-10/group | *$p < 0.05$<br>modified Bonferroni test |
| Figure 1B'<br>Cs-QSI | One way ANOVA for condition (Sham, HU, HU<sup>0.1G</sup>) $F_{[2,17]} = 2.2$; $p = 0.14$, | n=6/group | *$p < 0.05$<br>modified Bonferroni test |
| Figure 1C<br>104y>NaChBac Sleep | 3(Genotype) X 2(Sham, HU) X 24(Time) ANOVA $F_{[46,2024]} = 3.14$; $p = 1.29 \times 10^{-11}$ | n=12-16/group | |
| Figure 1D<br>104y>NaChBac STM | 3 (Genotype) X 2 (Sham, HU) ANOVA $F_{[2,39]} = 5.025$; $p = 0.01$. | n>=8/group | *$p < 0.05$<br>modified Bonferroni test |
| Figure 1D'<br>104y>NaChBac PSI | 3 (Genotype) X 2 (Sham, HU) ANOVA $F_{[2,27]} = 2.73$; $p = 0.083$ | n= 6/group | *$p < 0.05$<br>modified Bonferroni test |
| Figure 1D'<br>104y>NaChBac QSI | 3 (Genotype) X 2 (Sham, HU) ANOVA $F_{[2,27]} = 0.009$; $p = 0.991$ | n= 6/group | *$p < 0.05$<br>modified Bonferroni test |
| Figure G<br>Courtship Conditioning | One way ANOVA for condition (Sham, HU, HU<sup>0.1G</sup>) ANOVA $F_{[2,83]} = 9.089$; $p = 0.0003$ | n=28-29 flies per condition | *$p < 0.05$<br>modified Bonferroni test |
| Figure J<br>Cs-HU Olfactory learning | One way ANOVA for condition (Sham, HU, HU<sup>0.1G</sup>) $F_{[2,26]} = 15.03$; $p = 5.84 \times 10^{-5}$ | N=8-10/group | *$p < 0.05$<br>modified Bonferroni test |
|  | Statistics for Supplemental Figure S1 |  |  |
| Figure S1A<br>lio<sup>1</sup> STM | One way ANOVA for condition (veh, 0.1G, 0.1G<sup>SD</sup>) $F_{[2,26]} = 14.47$; $p = 5.95 \times 10^{-5}$ | n>=8/group | *$p < 0.05$<br>modified Bonferroni test |
| Figure S1B<br>lio<sup>1</sup> PSI | One way ANOVA for condition (veh, 0.1G, 0.1G<sup>SD</sup>) $F_{[2,17]} = 5.4$; $p = 0.01$ | n= 6/group | *$p < 0.05$<br>modified Bonferroni test |
| Figure S1B<br>*lio*<sup>1</sup> QSI | One way ANOVA for condition (veh, 0.1G, 0.1G<sup>SD</sup>) $F_{[2,18]} = 0.17$; $p=0.86$, n.s. $p>0.05$ | n= 5-8/group | *$p<0.05$<br>modified<br>Bonferroni<br>test |
| Figure S1C<br>*lio*<sup>2</sup> STM | One way ANOVA for condition (veh, 0.1G, 0.1G<sup>SD</sup>) $F_{[2,24]} = 7.60$; $p=0.002$; *$P<0.05$ | n=>8/group | *$p<0.05$<br>modified<br>Bonferroni<br>test |
| Figure S1D<br>*lio*<sup>2</sup> PSI | One way ANOVA for condition (veh, 0.1G, 0.1G<sup>SD</sup>) $F_{[2,13]} = 0.18$; $p=0.83$ | n=5-6/group | *$p<0.05$<br>modified<br>Bonferroni<br>test |
| Figure S1D<br>*lio*<sup>2</sup> QSI | One way ANOVA for condition (veh, 0.1G, 0.1G<sup>SD</sup>) $F_{[2,17]} = 2.66$; $p=0.09$ | n=5-10/group | *$p<0.05$<br>modified<br>Bonferroni<br>test |
| Figure S1F,G<br>*lio*<sup>2</sup> STM | paired t-test | n= 8-9/group | *$p<0.05$ |
| Figure S1H<br>*lat6* STM | One way ANOVA for condition (veh, 0.1G, 0.1G<sup>SD</sup>) $F_{[2,23]} = 5.48$; $p=0.01$ | n= 8-9/group | *$p<0.05$<br>modified<br>Bonferroni<br>test |
| Figure S1I<br>*lat6* PSI | One way ANOVA for condition (veh, 0.1G, 0.1G<sup>SD</sup>) $F_{[2,15]} = 1.98$; $p=0.17$ | n= 6/group | *$p<0.05$<br>modified<br>Bonferroni<br>test |
| Figure S1I<br>*lat6* QSI | One way ANOVA for condition (veh, 0.1G, 0.1G<sup>SD</sup>) $F_{[2,15]} = 1.57$; $p=0.23$. | n= 6/group | *$p<0.05$<br>modified<br>Bonferroni<br>test |
| Figure S1I<br>*elav>Orc3*<sup>RNAi</sup><br>STM | 3(genotype) X 2 (Drug) ANOVA $F_{[2,42]} = 8.5$; 0.001 | n= 8/group | *$p<0.05$<br>modified<br>Bonferroni<br>test |
| Figure S1K<br>*elav>Orc3*<sup>RNAi</sup><br>PSI | One way ANOVA for genotype $F_{[2,15]} = 2.8$; $p=0.09$ | n= 6/group | *$p<0.05$<br>modified<br>Bonferroni<br>test |
| Figure S1K<br>*elav>Orc3*<sup>RNAi</sup><br>QSI | One way ANOVA for genotype $F_{[2,14]} = 1.15$; $p=0.34$ | n=5/group | *$p<0.05$<br>modified<br>Bonferroni<br>test |
| Figure S1L<br>*CK2β*<sup>mbu/+</sup><br>STM | Student *t*-test *$p=0.0008$, | n=7-8 /group | |
| Figure S1N<br>247>RA<sup>CS</sup> | One way ANOVA for temperature (19°C, 30°C, 30°C + 0.1G) $F_{[2,40]} = 3.3$; $p=0.04$, | n = 10-17/group | *$p<0.05$<br>modified |
|  |  |  | Bonferroni test |
| Figure S1N<br>201y>RA<sup>CS</sup> | one way ANOVA for temperature (19°C, 30°C, 30°C + 0.1G)<br>F<sub>[2,25]</sub> = 2.2; p=0.13 , | n= 8-9/group | *p<0.05<br>modified<br>Bonferroni<br>test |

**Table S3.**
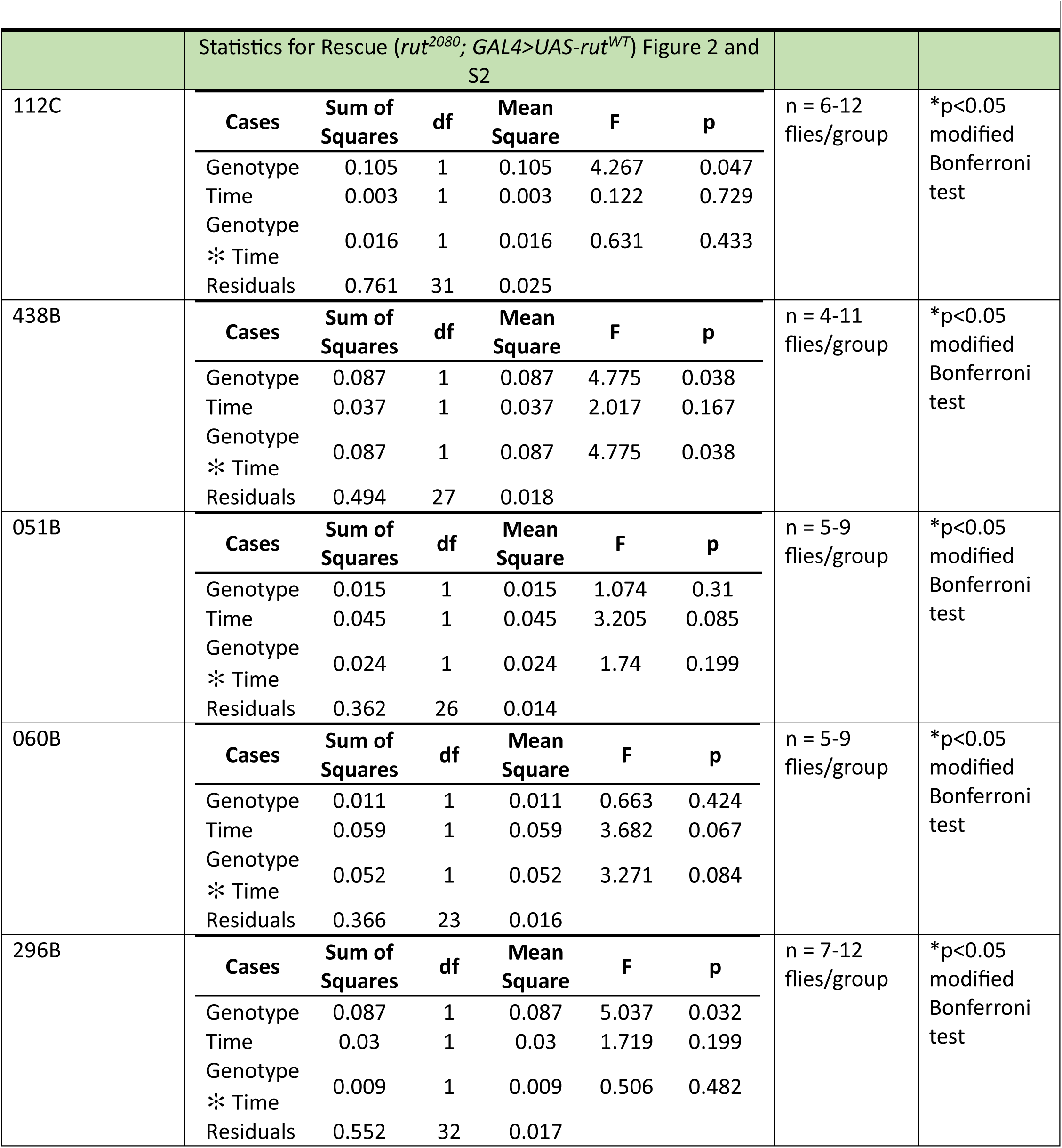
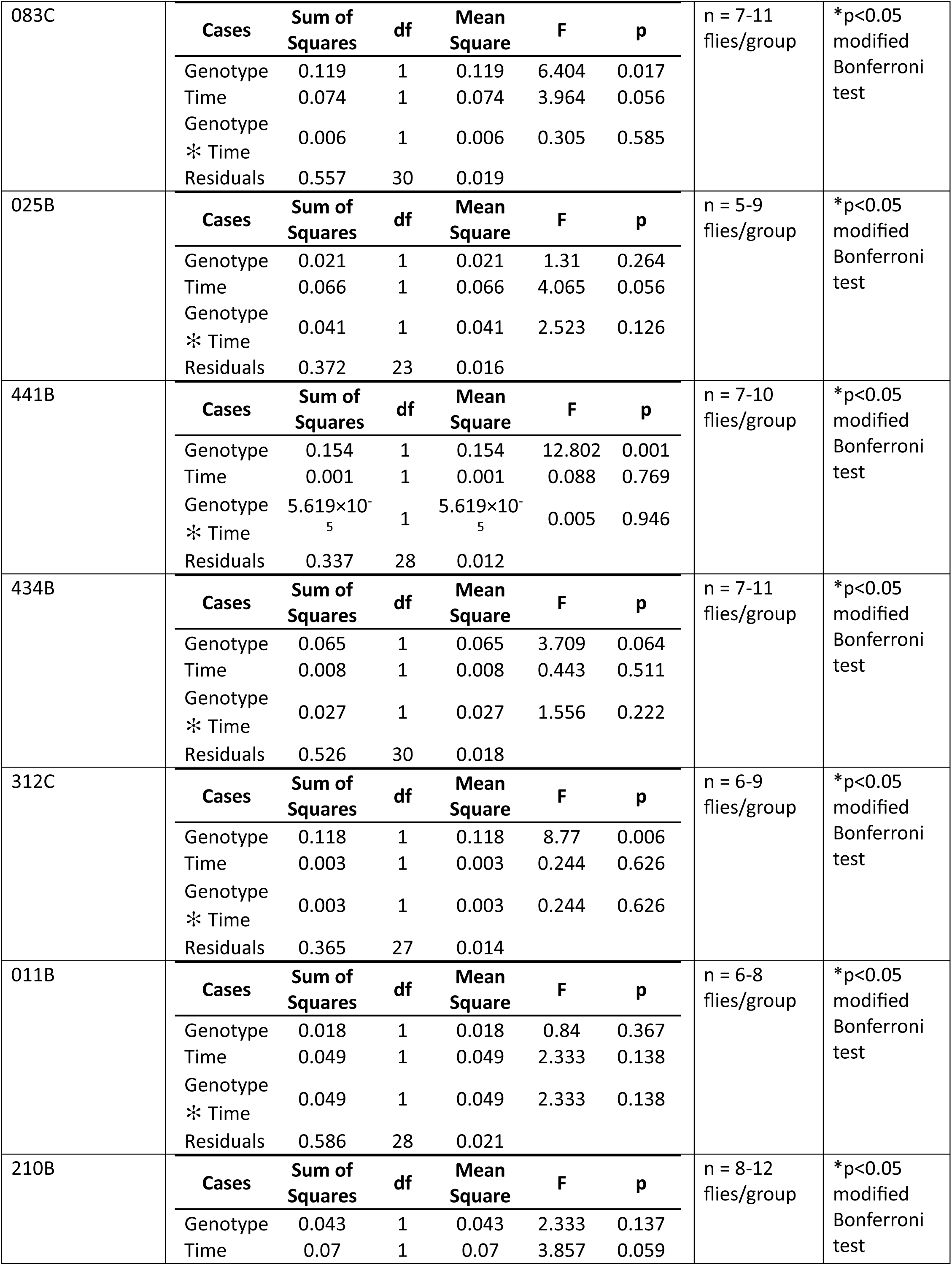
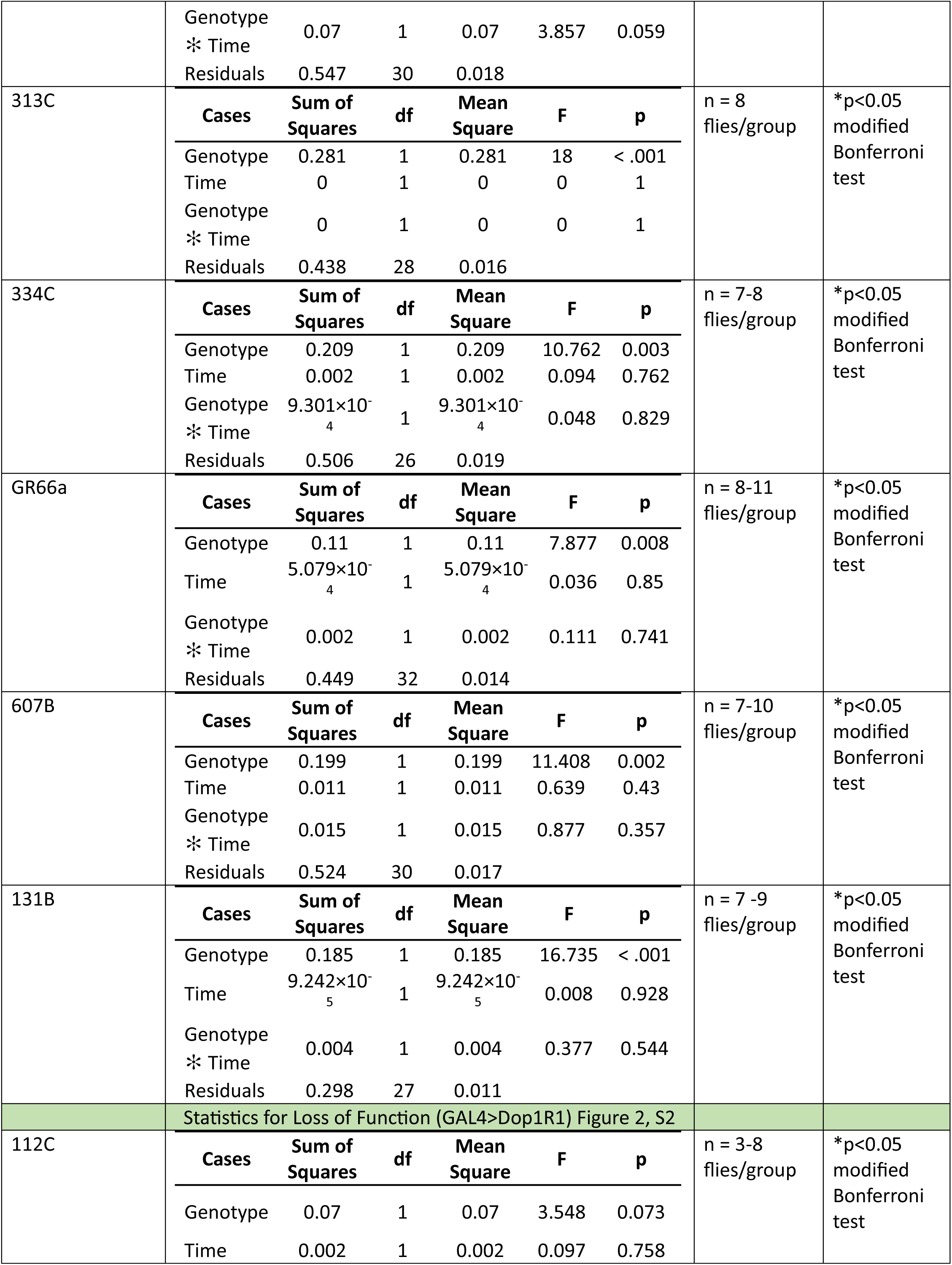
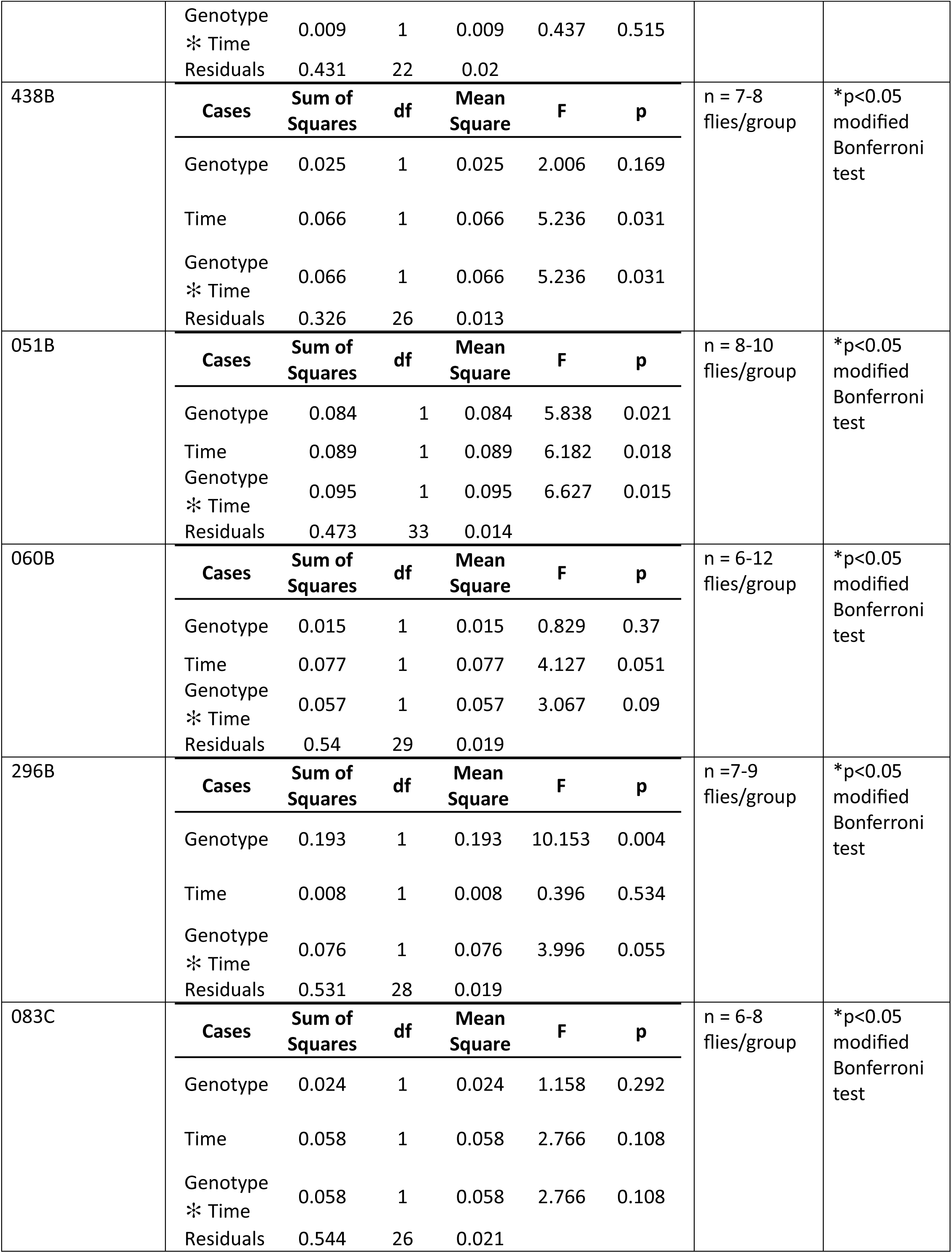
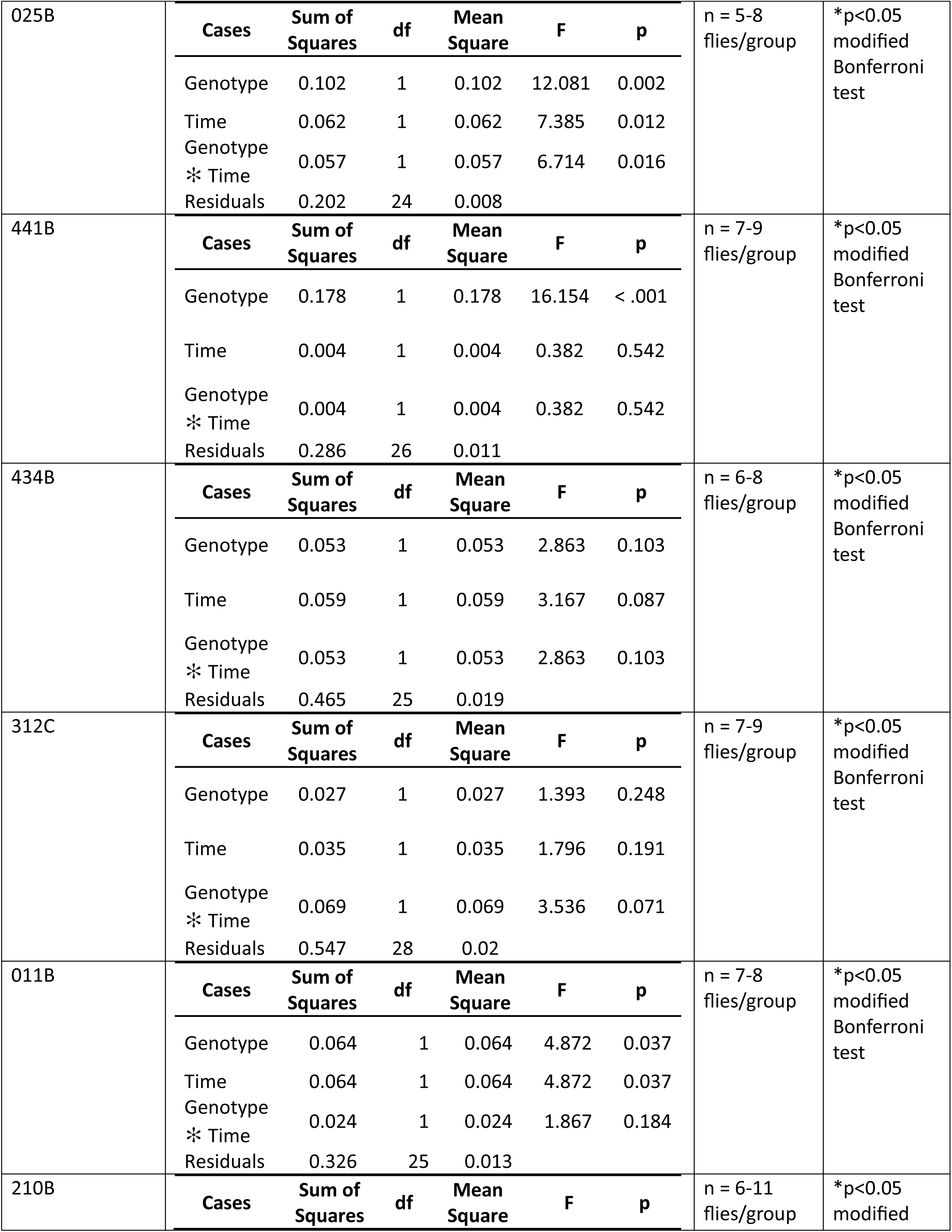
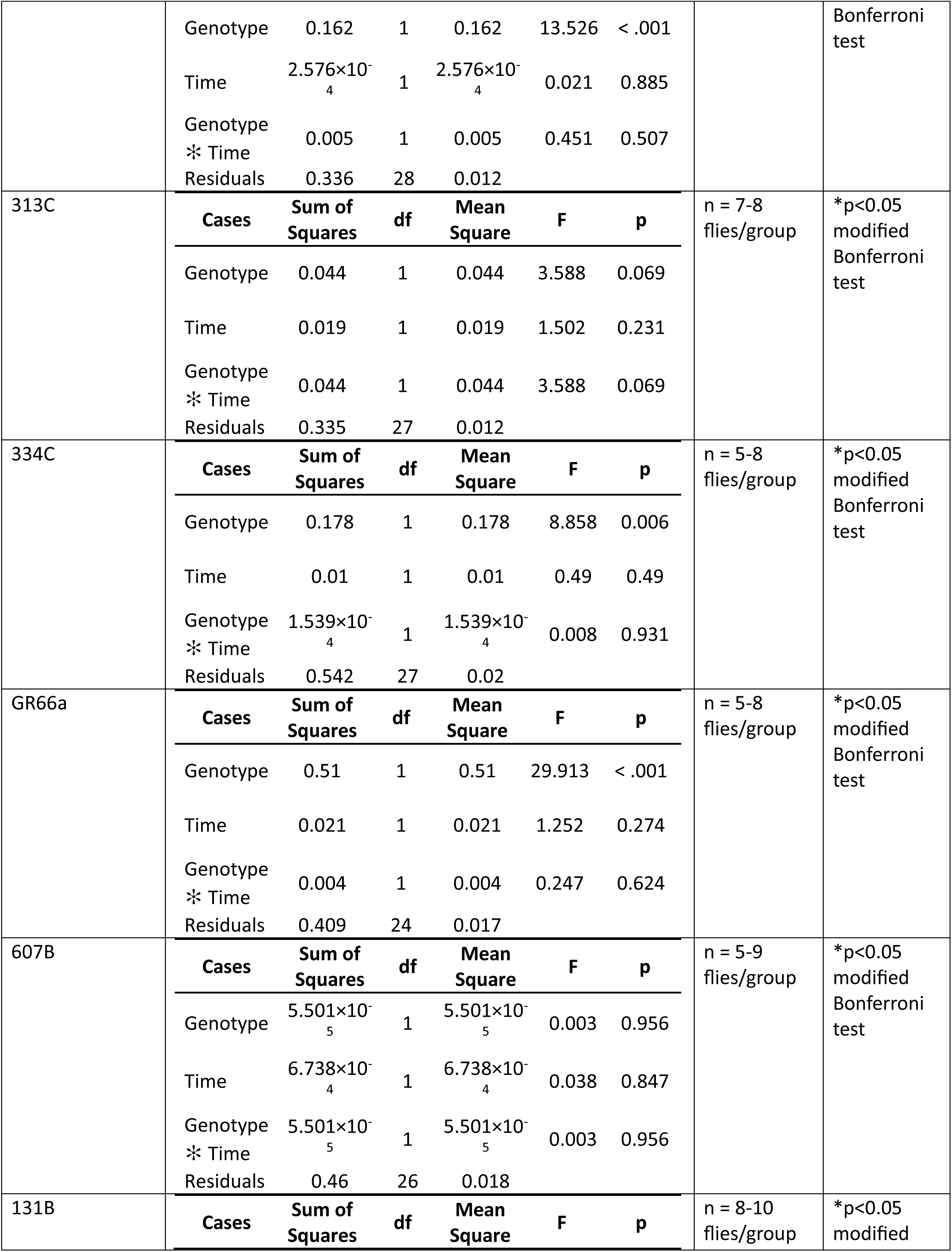
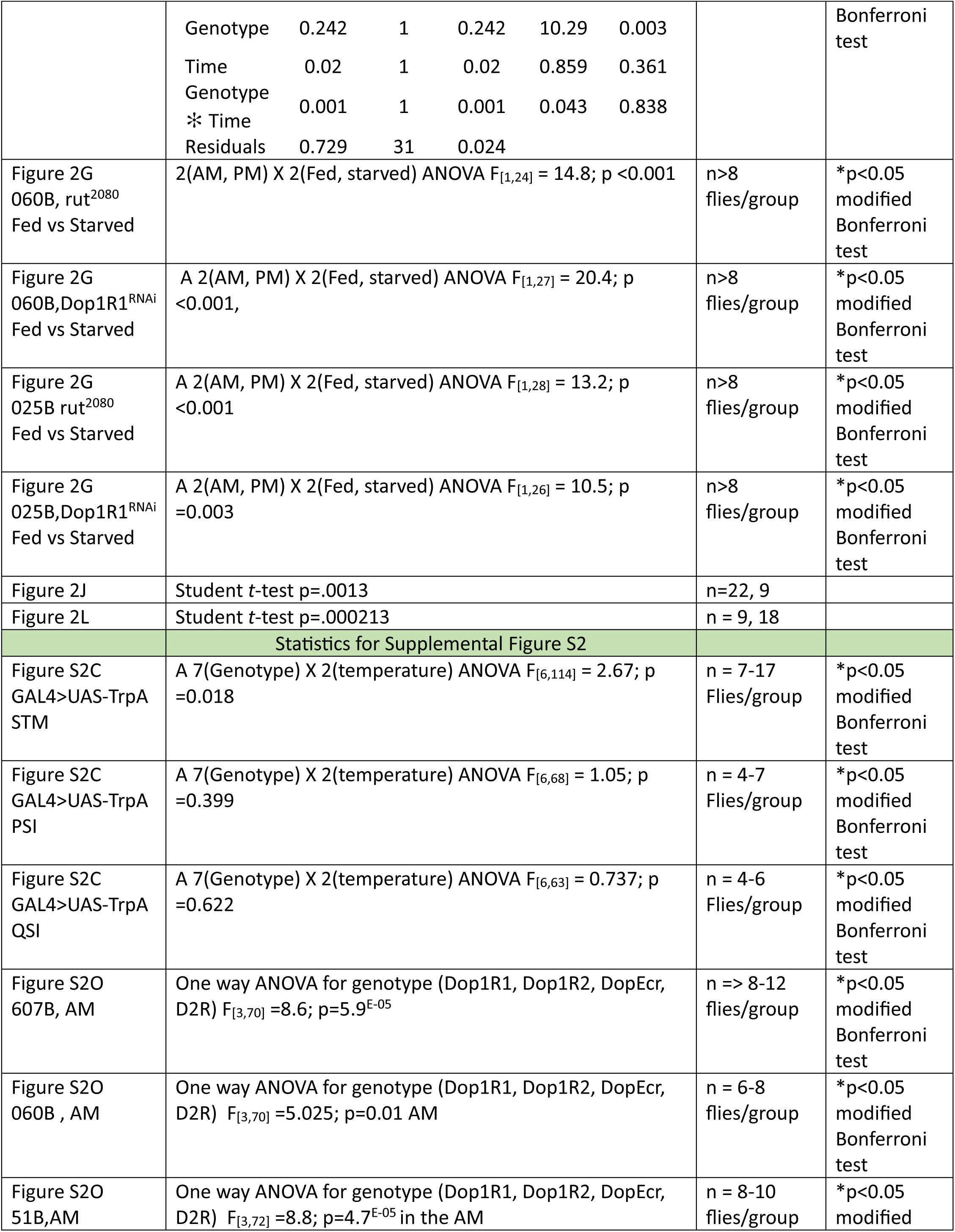
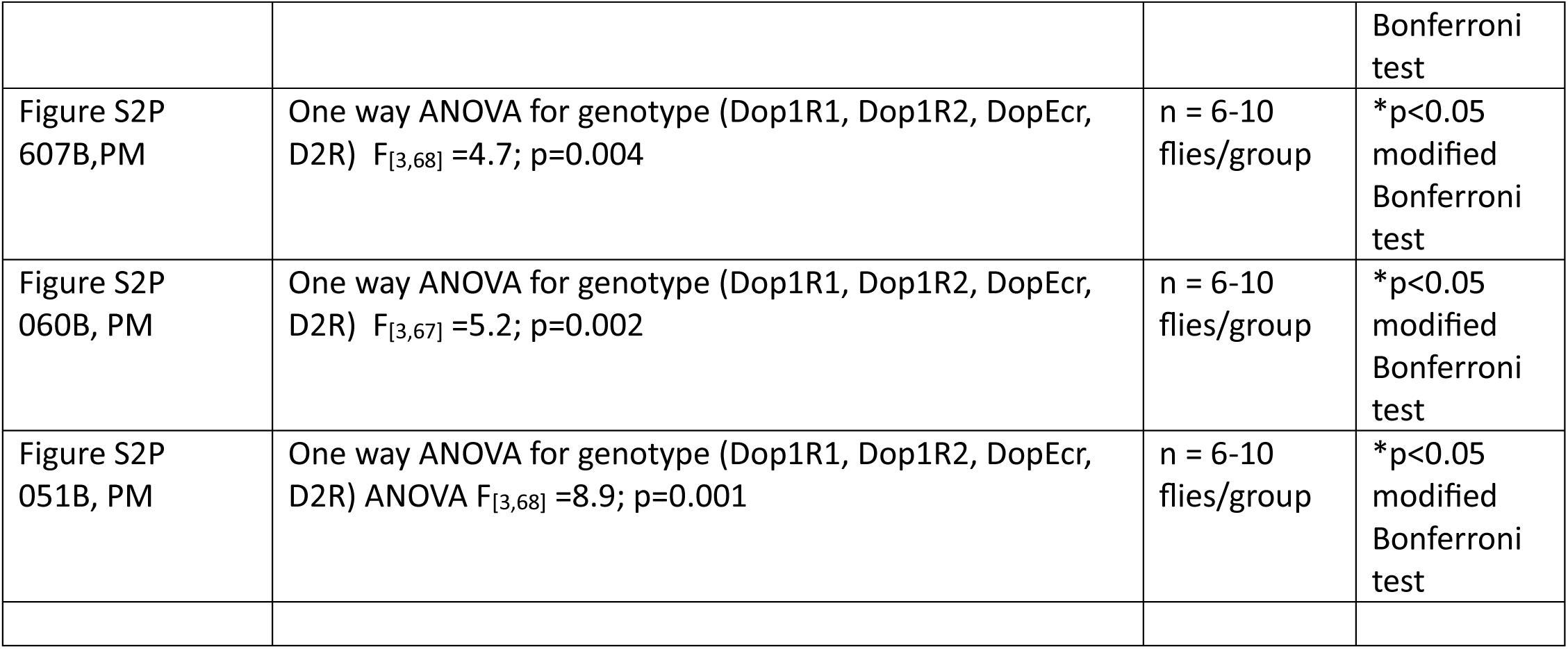

**Table S4.**
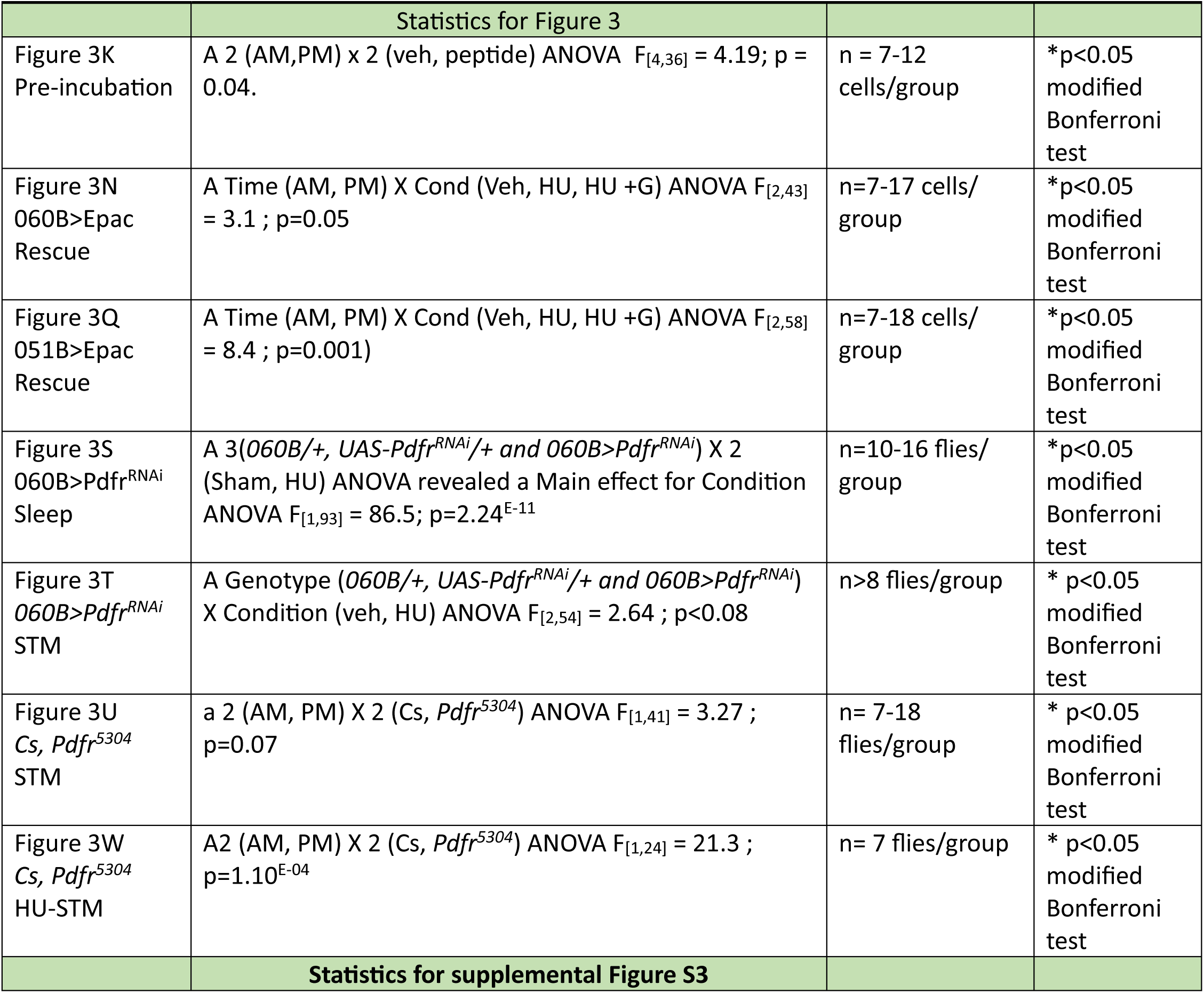
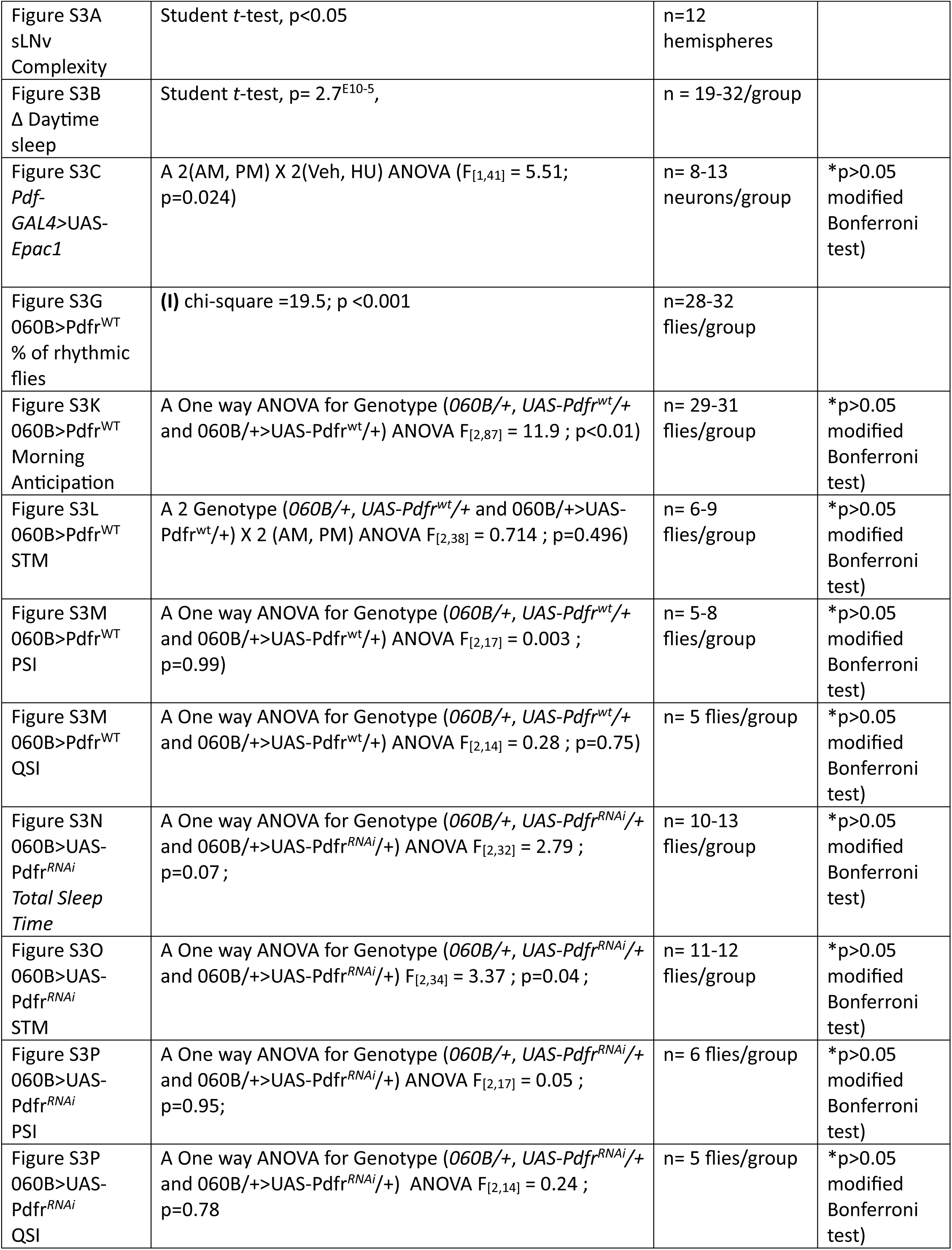
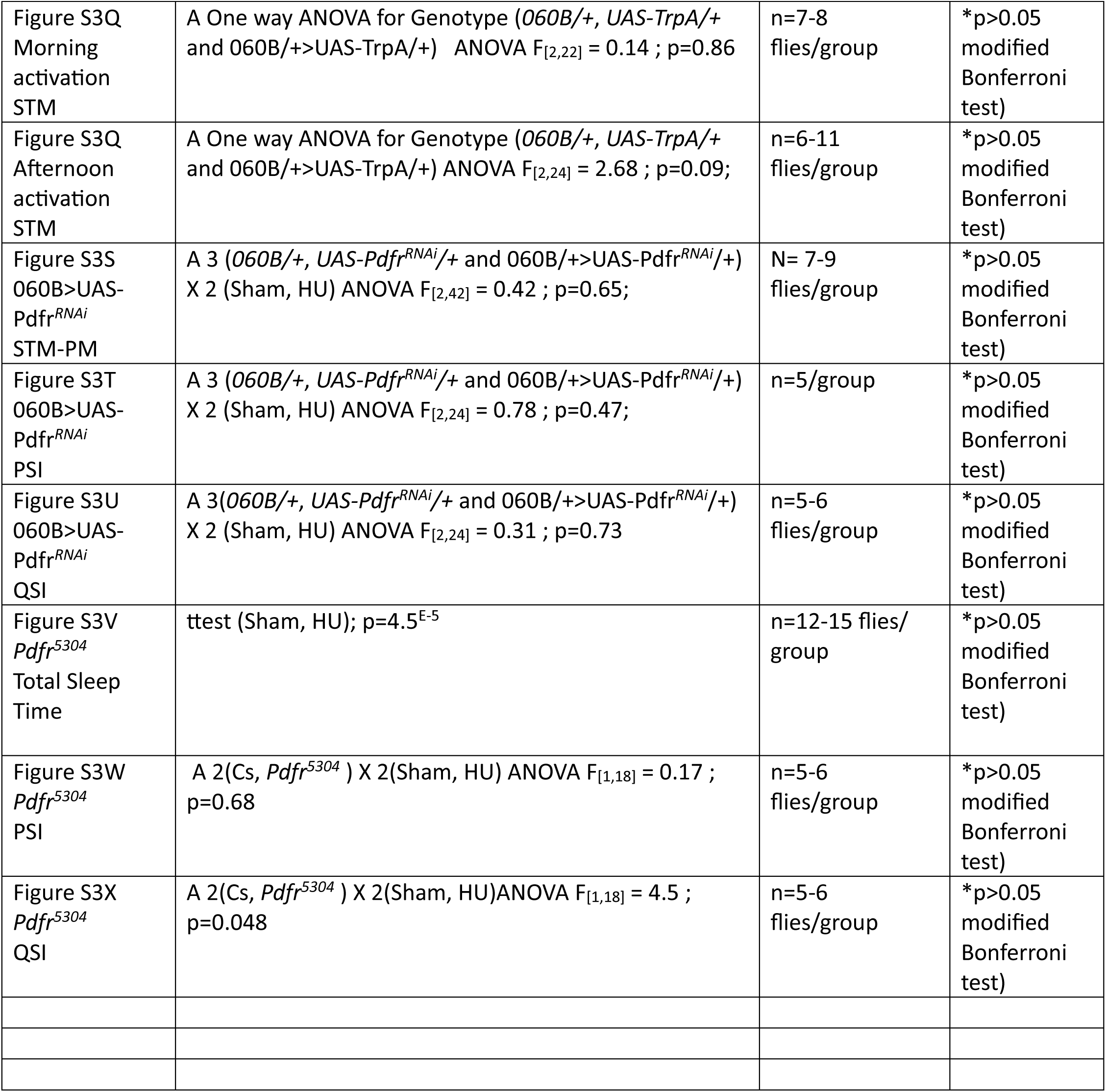

**Table S5.**
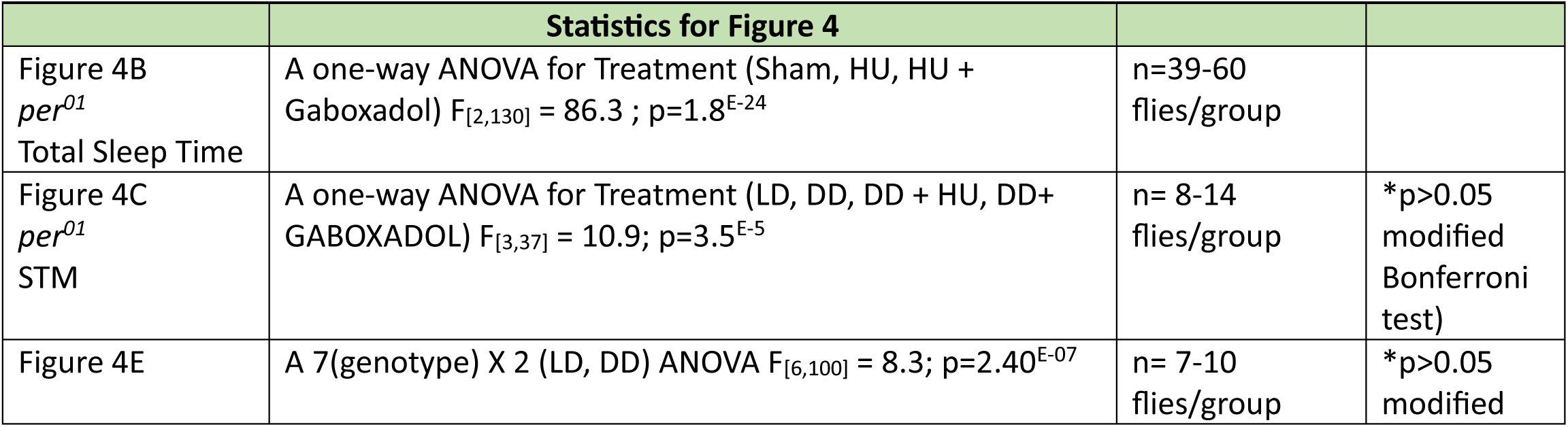
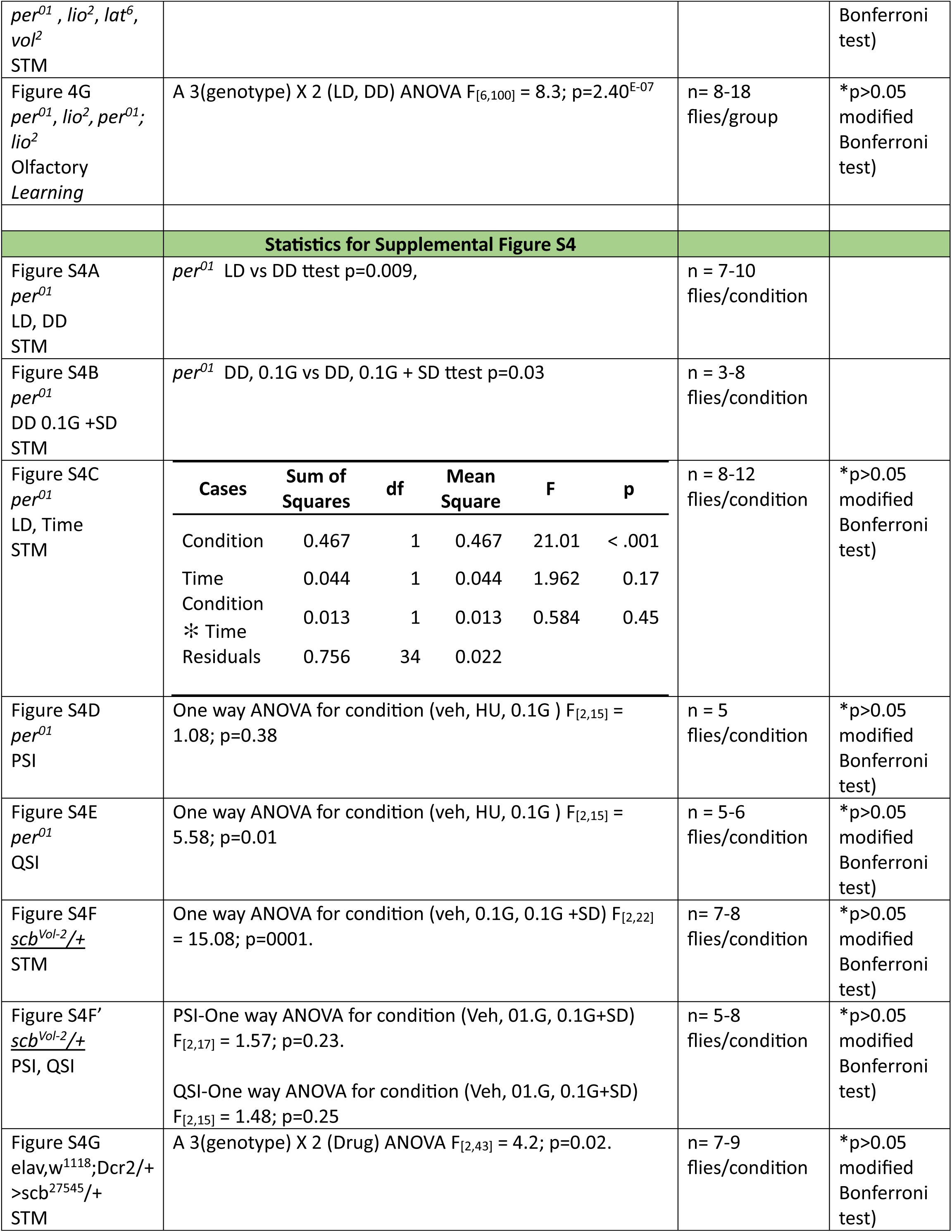
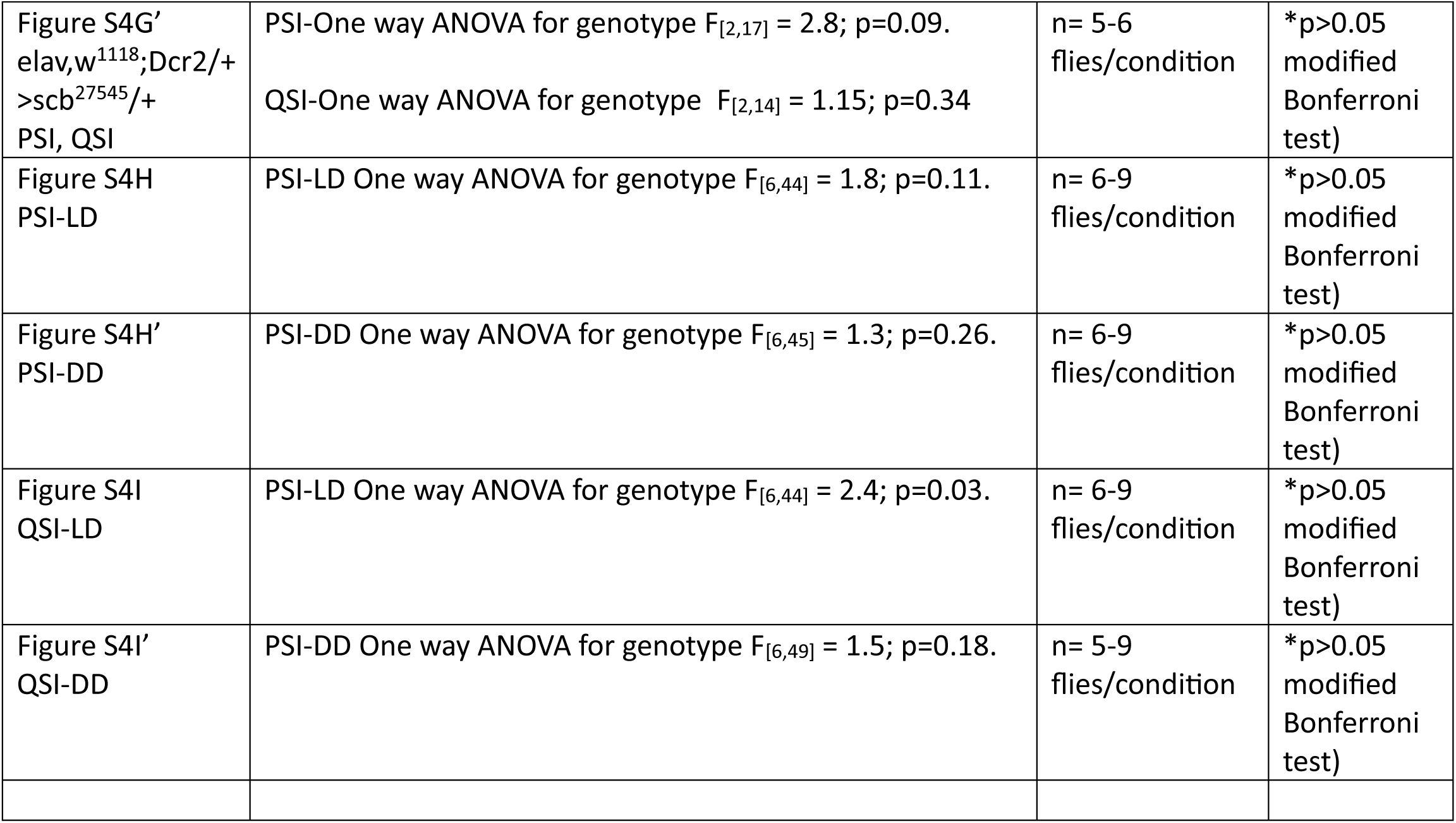

**Table S6.**

| Table S6 |  |  |  |  |  |  |  |
| --- | --- | --- | --- | --- | --- | --- | --- |
| Quantification of Figure 4D |  |  |  |  |  |  |  |
|  | *per*<sup>01</sup> | *lio*<sup>2</sup>/+ | *per*<sup>01</sup>; *lio*<sup>2</sup> | *lat*<sup>6</sup>/+ | *per*<sup>01</sup>; *lat*<sup>6</sup> | Vol<sup>2</sup>/+ | *per*<sup>01</sup>;Vol<sup>2</sup>/+ |
| Mean | 26.53 | 23.87 | 28.35 | 23.69 | 26.52 | 23.70 | 29.77 |
| SEM | 1.89 | 0.08 | 1.77 | 0.04 | 2.82 | 0.07 | 0.28 |
| % Rhythmic | 10.00% | 80.65% | 6.67% | 71.88% | 12.90% | 71.88% | 6.45% |
| n (rhythmic) | 3 | 25 | 2 | 23 | 4 | 23 | 2 |
| n (total) | 30 | 31 | 30 | 32 | 31 | 32 | 31 |

